# Human Pluripotent Stem Cell-Derived Neural Cells and Brain Organoids Reveal SARS-CoV-2 Neurotropism

**DOI:** 10.1101/2020.07.28.225151

**Authors:** Fadi Jacob, Sarshan R. Pather, Wei-Kai Huang, Samuel Zheng Hao Wong, Haowen Zhou, Feng Zhang, Beatrice Cubitt, Catherine Z. Chen, Miao Xu, Manisha Pradhan, Daniel Y. Zhang, Wei Zheng, Anne G. Bang, Hongjun Song, Juan Carlos de a Torre, Guo-li Ming

## Abstract

Neurological complications are common in patients with COVID-19. While SARS-CoV-2, the causal pathogen of COVID-19, has been detected in some patient brains, its ability to infect brain cells and impact their function are not well understood, and experimental models using human brain cells are urgently needed. Here we investigated the susceptibility of human induced pluripotent stem cell (hiPSC)-derived monolayer brain cells and region-specific brain organoids to SARS-CoV-2 infection. We found modest numbers of infected neurons and astrocytes, but greater infection of choroid plexus epithelial cells. We optimized a protocol to generate choroid plexus organoids from hiPSCs, which revealed productive SARS-CoV-2 infection that leads to increased cell death and transcriptional dysregulation indicative of an inflammatory response and cellular function deficits. Together, our results provide evidence for SARS-CoV-2 neurotropism and support use of hiPSC-derived brain organoids as a platform to investigate the cellular susceptibility, disease mechanisms, and treatment strategies for SARS-CoV-2 infection.

## INTRODUCTION

Severe acute respiratory syndrome coronavirus 2 (SARS-CoV-2), the pathogen responsible for the COVID-19 pandemic, has infected over 15 million people and has contributed to over 60,000 deaths world-wide within this year so far (WHO, 2020). While the disease primarily affects the respiratory system, damage and dysfunction has also been found in other organs, including the kidney, heart, liver, and brain (Yang et al., 2020b). Neurological complications, such as cerebrovascular injury, altered mental status, encephalitis, encephalopathy, dizziness, headache, hypogeusia, and hyposmia, as well as neuropsychiatric ailments, including new onset psychosis, neurocognitive syndrome, and affective disorders, have been reported in a significant number of patients (Mao et al., 2020; Varatharaj et al., 2020). Viral RNA has been detected in the brain and cerebrospinal fluid (CSF) of some patients with COVID-19 and concomitant neurological symptoms (Helms et al., 2020; Moriguchi et al., 2020; Puelles et al., 2020; Solomon et al., 2020). Despite numerous reports of neurological findings in patients with COVID-19, it remains unclear whether these symptoms are a consequence of direct neural infection, para-infectious or post-infectious immune-mediated disease, or sequalae of systemic disease (Ellul et al., 2020). In addition, variability in patient presentation and inconsistent timing of testing for viral RNA in the CSF or brain limit the interpretation of results in human patients. Limited availability of post-mortem brain tissue from patients with COVID-19 and the inability to study ongoing disease pathogenesis further underscore the need for an accessible and tractable experimental model to investigate SARS-CoV-2 neurotropism, its functional impact and for future therapeutic treatment testing.

Classic animal models, such as rodents, are limited in their ability to recapitulate human COVID-19 symptoms and usually require viral or transgenic mediated overexpression of human SARS-CoV-2 receptor angiotensin-converting enzyme 2 (ACE2) to yield symptoms (Bao et al., 2020; Sun et al., 2020). Although cell lines, including many human cancer cell lines, have been used to study SARS-CoV-2 infection and test drug efficacy (Hoffmann et al., 2020b; Ou et al., 2020; Shang et al., 2020; Wang et al., 2020), they do not accurately recapitulate normal human cell behavior and often harbor tumor-associated mutations, such as TP53, which may affect SARS-CoV-2 replication or the cellular response to SARS-CoV-2 infection (Ma-Lauer et al., 2016). Furthermore, human tissue and organ systems contain multiple cell types with variable levels of ACE2 expression and viral susceptibility, which are not adequately represented in these human cell lines. These limitations support the development of a human cellular model for SARS-CoV-2 infection that better recapitulates the cellular heterogeneity and function of individual tissues.

Human pluripotent stem cell (hiPSC)-based models provide an opportunity to investigate the susceptibility of various brain cell types to viral infection and their consequences. hiPSCs have been used to generate a variety of monolayer and three dimensional (3D) organoid cultures to study human diseases and potential treatments. For example, hiPSC-derived neural progenitors in monolayer and brain organoids were instrumental in studying the impact of Zika virus (ZIKV) infection on human brain development and solidifying the link between ZIKV infection of neural progenitor cells and microcephaly seen in newborns (Ming et al., 2016; Qian et al., 2016; Tang et al., 2016). Additionally, these cultures were useful in screening for drugs to treat ZIKV infection (Xu et al., 2016). Recently, hiPSC-derived organoids have been used to model SARS-CoV-2 infection in many organs, including the intestine, lung, kidney, liver, pancreas, and vasculature (Lamers et al., 2020; Monteil et al., 2020; Yang et al., 2020a; Zhou et al., 2020). These studies have shown that SARS-CoV-2 can infect and replicate within cells of multiple organs, leading to transcriptional changes indicative of inflammatory responses and altered cellular functions. Here, we used hiPSC-derived neurons, astrocytes, and microglia in monolayer cultures and region-specific brain organoids of the cerebral cortex, hippocampus, hypothalamus, and midbrain to investigate the susceptibility of brain cells to SARS-CoV-2 infection. We observed modest numbers of infected neurons and astrocytes, with the exception of regions of organoids with choroid plexus epithelial cells that exhibited high levels of infectivity. Using an optimized protocol to generate choroid plexus organoids (CPOs) from hiPSCs, we showed evidence of productive infection by SARS-CoV-2 and further investigated the functional consequences at the cellular and molecular levels.

## RESULTS

### SARS-CoV-2 Neurotropism in Various hiPSC-derived Brain Cells and Organoids

To investigate the susceptibility of human brain cells to SARS-CoV-2 infection, we tested various hiPSC-derived neural cells in monolayer cultures and region-specific brain organoids generated using several established (Qian et al., 2018; Qian et al., 2016) and modified protocols (Sakaguchi et al., 2015) (Figure 1A). Monolayer hiPSC-derived cortical neurons, microglia, and astrocytes were exposed to either SARS-CoV-2 virus isolate or vehicle control for 12 hours and analyzed at 24, 48, and 120 hours post-infection (hpi) for immunolabelling using convalescent serum from a patient with COVID-19 or SARS-CoV-2 nucleoprotein antibodies (Figures S1A-C). We confirmed the identity of neurons, microglia, and astrocytes by immunostaining for markers MAP2, PU.1, and GFAP, respectively (Figures S1A-C). hiPSC-derived cortical neurons were co-cultured on hiPSC-derived astrocytes to improve survival, and analysis at 120 hpi showed rare infection of MAP2^+^ cells (Figure S1A). At 48 and 120 hpi, hiPSC-derived microglia showed no infection of PU.1^+^ cells (Figure S1B), whereas hiPSC-derived astrocytes showed sparse infection of GFAP^+^ cells (Figure S1C). As a validation, we tested infection of primary human astrocytes and observed sparse infection at 24, 48, and 120 hpi (Figure S1D). Quantification of hiPSC-derived and primary human astrocytes showed infection of 0.02% and 0.18% of all cells at 120 hpi, respectively (Figure S1E). These results revealed the ability of SARS-CoV-2 to infect monolayer human cortical neurons and astrocytes, but not microglia, although infection of neurons and astrocytes was rare.

**Figure 1.**
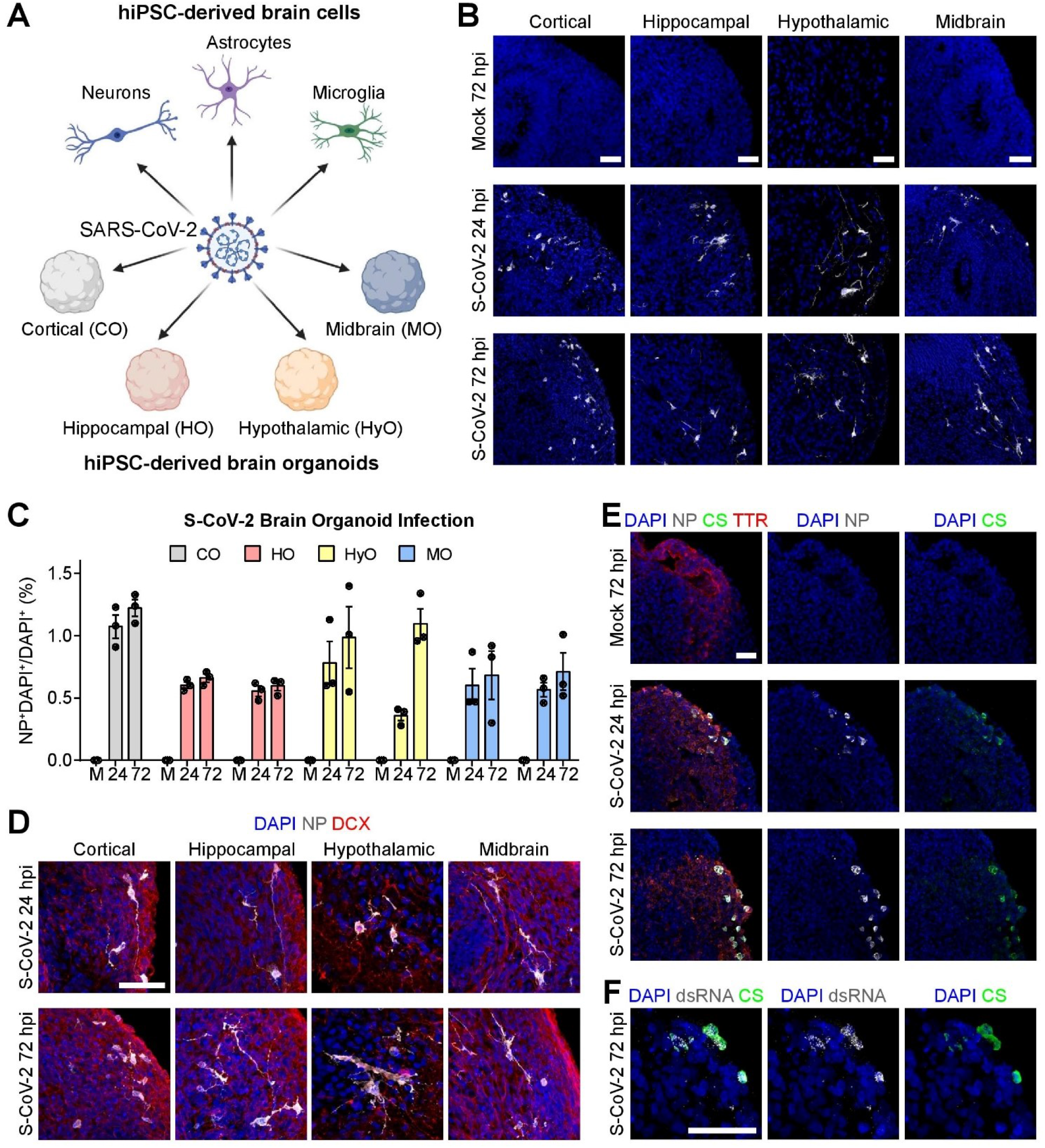
SARS-CoV-2 Neurotropism in hiPSC-derived Brain Organoids. **(A)** Diagram outlining the strategy for testing the broad susceptibility of brain cells to SARS-CoV-2 infection. Tested cultures include hiPSC-derived cortical neurons, astrocytes, and microglia in monolayer cultures, and hiPSC-derived cortical, hippocampal, hypothalamic, and midbrain organoids. **(B)** Representative confocal images of fluorescent immunohistology for DAPI and SARS-CoV-2 nucleoprotein (NP) in hiPSC-derived cortical, hippocampal, hypothalamic, and midbrain organoids after SARS-CoV-2 (S-CoV-2) or vehicle treatment at 24 and 72 hours post-infection (hpi). Scale bars, 50 μm. **(C)** Quantification of percentages of NP^+^DAPI^+^/DAPI^+^ cells in S-CoV-2 and vehicle treated hiPSC-derived cortical (CO), hippocampal (HO), hypothalamic (HyO), and midbrain (MO) organoid cultures at 24 and 72 hpi. Values represent mean ± SEM with individual data points plotted (n = 3 organoids per brain region and condition with 3 images per organoid). Brain organoids derived from two independent hiPSC lines were analyzed for HO, HyO and MO. **(D)** Representative confocal images of fluorescent immunohistology for DAPI, NP, and neuronal marker doublecortin (DCX) in hiPSC-derived cortical, hippocampal, hypothalamic, and midbrain organoids after S-CoV-2 or vehicle treatment at 24 and 72 hpi. Scale bar, 50 μm. **(E)** Representative confocal images of fluorescent immunohistology for DAPI, NP, convalescent serum from a patient with COVID-19 (CS), and transthyretin (TTR) in hippocampal organoids after S-CoV-2 or vehicle treatment at 24 and 72 hpi highlighting regions with choroid plexus cell differentiation. Scale bar, 50 μm. **(F)** Representative confocal images of fluorescent immunohistology for DAPI, double-stranded RNA (dsRNA), and patient convalescent serum (CS) in a hippocampal organoid with a region of choroid plexus differentiation after S-CoV-2 treatment at 72 hpi. Scale bar, 50 μm. Also see **Figure S1**.

To examine the neurotropism of SARS-CoV-2 in a model system that more closely resembles the human brain, we exposed cortical, hippocampal, hypothalamic, and midbrain organoids to SARS-CoV-2 or vehicle control for 8 hours and analyzed samples at 24 and 72 hpi. We confirmed the regional identity of cortical, hippocampal, hypothalamic, and midbrain organoids by immunostaining with the markers CTIP2, PROX1, OTP, and TH, respectively (Figure S1F). SARS-CoV-2 nucleoprotein was detected sparsely in these organoids derived from two different hiPSC lines in a range that averaged between 0.6% and 1.2% of all cells at 24 and 72 hpi (Figures 1B, 1C, and S1G). Co-immunolabeling with DCX and SARS-CoV-2 nucleoprotein identified most of the infected cells as neurons in all organoids (Figure 1D) and infection of some GFAP^+^ astrocytes was found in hypothalamic organoids (Figure S1H). Because most of the cells within these brain organoids were neurons, the relative susceptibility of neurons compared to astrocytes could not be assessed with certainty. The number of infected cells did not significantly increase from 24 to 72 hpi (Figures 1C), indicating that infection may not spread among neurons in brain organoids.

During development, the choroid plexus develops adjacent to the hippocampus (Lun et al., 2015) and some of our hippocampal organoids contained regions with choroid plexus epithelial cells, which were identified by transthyretin (TTR) expression (Figure 1E). Notably, we observed a greater density of infected cells in these regions (Figure 1E). Additionally, we observed co-localization of patient convalescent serum and double-stranded RNA (dsRNA) immunolabeling in regions with choroid plexus cells, further supporting SARS-CoV-2 infection (Figure 1F).

Together, these results demonstrate that SARS-CoV-2 exhibits modest tropism for neurons and astrocytes of multiple brain regions, but higher infectivity of choroid plexus epithelial cells.

### Generation of Choroid Plexus Organoids from hiPSCs

To validate the higher susceptibility and further investigate consequences of SARS-CoV-2 infection of choroid plexus cells, we sought to generate more pure choroid plexus organoids from hiPSCs. The most dorsal structures of the telencephalon, including the choroid plexus, are patterned by high WNT and BMP signaling from the roof plate (Lun et al., 2015). An early *in vitro* study demonstrated the sufficiency of BMP4 exposure to induce choroid plexus from neuroepithelial cells (Watanabe et al., 2012). Furthermore, exposure of human embryonic stem cell-derived embryoid bodies to the GSK3β antagonist CHIR-99021 and BMP4 was shown to generate 3D choroid plexus tissue (Sakaguchi et al., 2015). Although these studies were instrumental in understanding choroid plexus specification from stem cells, they are limited by variable efficiency and have uncertain applicability to hiPSCs. Building upon these pioneering studies, we optimized a simple protocol to generate choroid plexus organoids (CPOs) from hiPSCs (Figure 2A). Undifferentiated hiPSCs grown in a feeder-free condition were aggregated into embryoid bodies consisting of approximately 5,000 cells each using an Aggrewell plate (Figure 2A). Embryoid bodies were patterned to anterior neuroectodermal fate using dual-SMAD inhibition combined with WNT inhibition (Figure 2A). At 8 days *in vitro* (DIV), neural progenitors were patterned towards the choroid plexus fate by promoting high WNT signaling using the GSK3β antagonist CHIR-99021 and high levels of human recombinant BMP-7. CPOs maintained a round morphology at 15 DIV and they expressed medial forebrain markers LMX1A and OTX2, with minimal numbers of FOXG1^+^ cells at 20 DIV, indicating choroid plexus progenitor fate (Figures 2B, 2C and S2A). CPOs began to form more translucent cellular extensions by 25 DIV that produce thinner projections lined by cuboidal cells by 50 DIV (Figure 2A). At 50 DIV, CPOs displayed morphology resembling the human choroid plexus epithelium with extensions of cuboidal epithelial cells expressing choroid plexus markers OTX2, Aquaporin 1 (AQP1), and TTR (Figures 2D and S2B). Quantification of cells expressing various markers showed very high purity and consistency across two hiPSC lines (Figures 2C and 2E).

**Figure 2.**
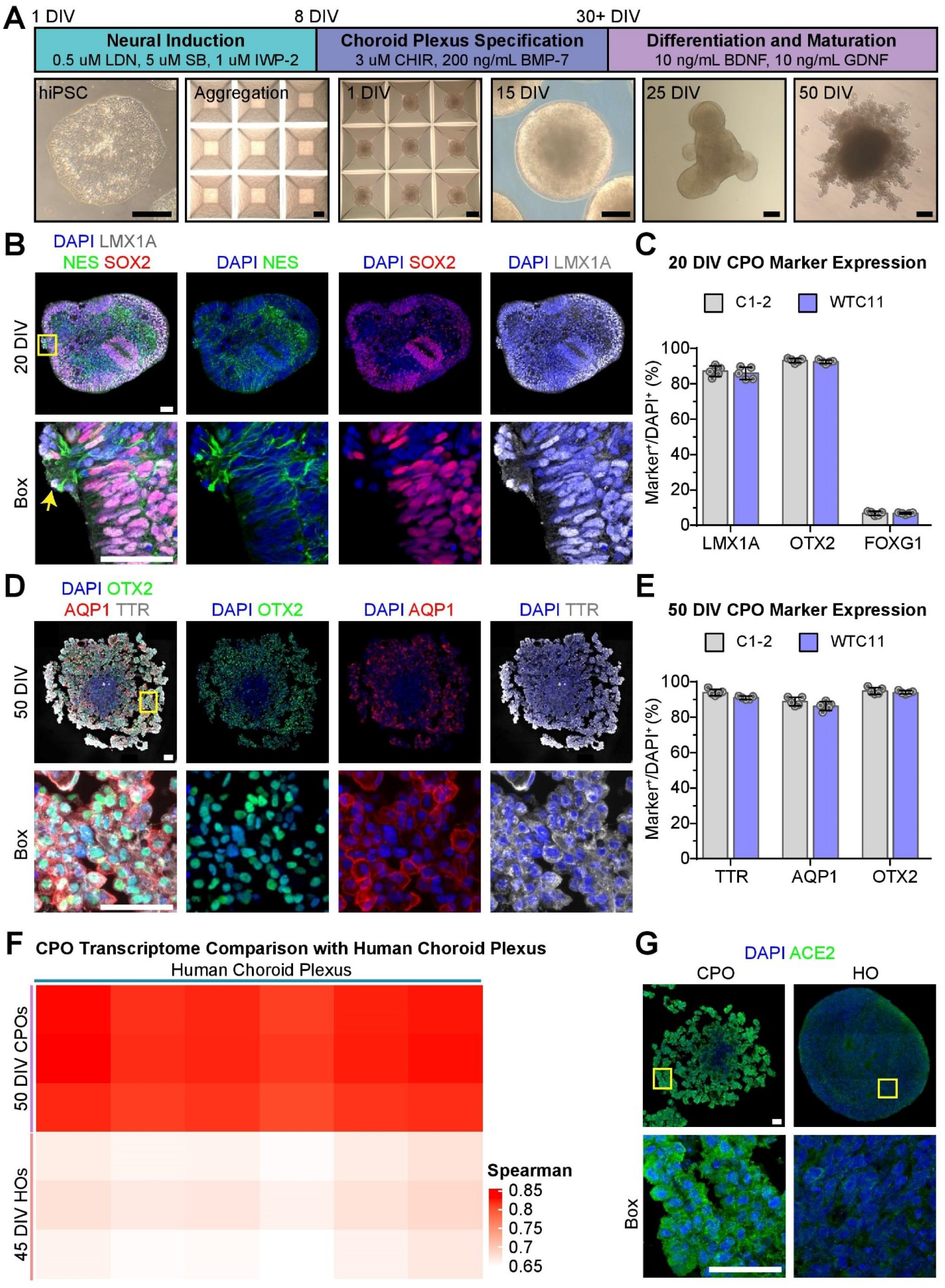
Generation of Choroid Plexus Organoids from hiPSCs. **(A)** Diagram describing the protocol for generating choroid plexus organoids (CPOs) from hiPSCs with sample bright-field images of hiPSCs, aggregated hiPSCs, and CPOs at 1, 15, 25, and 50 days *in vitro* (DIV). Scale bars, 200 μm. **(B)** Representative confocal images of fluorescent immunohistology for DAPI, LMX1A, NESTIN (NES), and SOX2 in CPOs at 20 DIV. Yellow arrow highlights a region beginning to differentiate into choroid plexus from the neuroepithelium. Scale bars, 50 μm. **(C)** Quantification of percentages of LMX1A^+^/DAPI^+^, OTX2^+^/DAPI^+^, and FOXG1^+^/DAPI^+^ cells in CPOs at 20 DIV. Values represent mean ± SEM with individual data points plotted (n = 5 organoids per hiPSC line with 3 images per organoid). **(D)** Representative confocal images of fluorescent immunohistology for DAPI, OTX2, AQP1, and TTR in CPOs at 50 DIV. Scale bars, 50 μm. **(E)** Quantification of percentages of TTR^+^/DAPI^+^, AQP1^+^/DAPI^+^, and OTX2^+^/DAPI^+^ cells in CPOs at 50 DIV. Values represent mean ± SEM with individual data points plotted (n = 5 organoids per hiPSC line with 3 images per organoid). **(F)** Heatmap comparing the Spearman correlation of the bulk RNA transcriptomes of 50 DIV CPOs and 45 DIV hippocampal organoids (HOs) to adult human choroid plexus tissue (Rodriguez-Lorenzo et al., 2020). **(G)** Representative confocal images of fluorescent immunohistology for DAPI and ACE2 in the CPO and HO. Scale bars, 50 μm. Also see **Figure S2**.

To further characterize these CPOs, we performed transcriptome analysis of CPOs at 50 DIV by bulk RNA sequencing (RNA-seq). The transcriptome of CPOs, but not that of hippocampal organoids at 45 DIV, showed high correlation to published transcriptomes of the adult human choroid plexus tissue (Rodriguez-Lorenzo et al., 2020) (Figure 2F). Detailed analysis further showed that CPOs, but not hippocampal organoids, expressed multiple markers for choroid plexus epithelial cells at similar levels as adult human choroid plexus tissue, including TTR, AQP1, Chloride Intracellular Channel 6 (CLIC6), Keratin 18 (KRT18), MSX1, and LMX1A (Figure S2C).

Given our initial finding of a high rate of infection of choroid plexus cells by SARS-CoV-2, we examined the expression of known receptors for SARS-CoV-2. Immunohistology showed expression of the key SARS-CoV-2 receptor ACE2 at a much higher level in CPOs than in hippocampal organoids (Figure 2G). RNA-seq analysis also showed expression of ACE2 and TMPRSS2 in CPOs at levels similar to adult human choroid plexus tissue, but much higher than in hippocampal organoids (Figure S2D). Expression levels of NRP1, a newly identified receptor for SARS-CoV-2 (Cantuti-Castelvetri et al., 2020), were similar in all three samples.

Together, these results show that our CPOs exhibit a similar transcriptome as adult human choroid plexus tissue and express markers for choroid plexus epithelial cells and SARS-CoV-2 receptors, representing a suitable experimental model to study SARS-CoV-2 infection.

### Productive Infection of hiPSC-derived Choroid Plexus Organoids by SARS-CoV-2 and Cellular Consequences

Next, we exposed CPOs at 47 DIV to either SARS-CoV-2 or vehicle control for 8 hours and analyzed samples at 24 and 72 hpi. Upon SARS-CoV-2, but not vehicle treatment, many TTR^+^ choroid plexus cells showed infection as identified by co-localization of patient convalescent serum and SARS-CoV-2 nucleoprotein immunolabeling in CPOs derived from two independent hiPSC lines (Figures 3A and S3A). Additionally, we observed co-localization of patient convalescent serum and abundant dsRNA immunolabeling in TTR^+^ choroid plexus cells (Figure S3B). Infected cells identified by SARS-CoV-2 nucleoprotein immunostaining also showed high levels of ACE2 expression, consistent with ACE2 being a critical cell entry receptor for SARS-CoV-2 (Figure S3C). Quantification revealed that an average of 9.0% of TTR^+^ cells were infected at 24 hpi across two hiPSC lines (Figure 3B). The percentage of infected cells increased significantly from 24 to 72 hpi in both hiPSC lines, suggesting that infection could be productive and spread among cells (Figure 3B). To confirm productive infection, we examined viral titer using culture supernatants and CPO lysates at 0, 24, and 72 hpi. Indeed, there was a significant increase in titers of infectious virus over time after the initial infection (Figure 3C). The increase of infectious virus present in CPO lysates superseded the increase in culture supernatants, suggesting that intracellular viral production by 24 hpi is not significantly released until later, which coincides with our observation of an increased number of infected cells at 72 hpi (Figures 3B and 3C).

**Figure 3.**
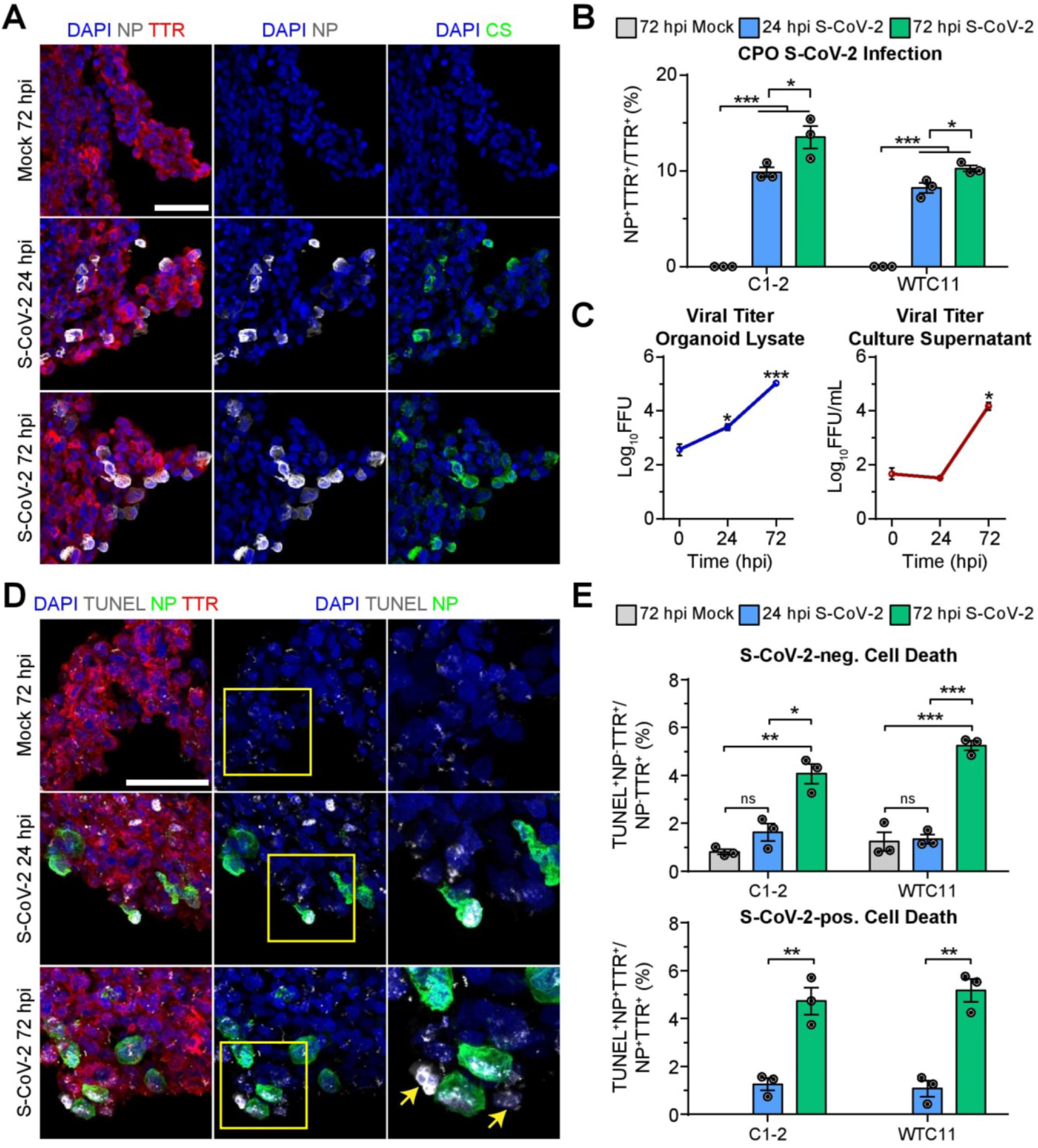
Productive Infection of Choroid Plexus Organoids and Increased Cell Death by SARS-CoV-2. **(A)** Representative confocal images of fluorescent immunohistology for DAPI, SARS-CoV-2 nucleoprotein (NP), patient convalescent serum (CS), and transthyretin (TTR) in CPOs after S-CoV-2 or vehicle treatment at 24 and 72 hpi. Scale bar, 50 μm. **(B)** Quantification of percentages of NP^+^TTR^+^/TTR^+^ cells in CPOs after S-CoV-2 or vehicle treatment at 24 and 72 hpi. Values represent mean ± SEM with individual data points plotted (n = 3 organoids per hiPSC line with 3 images per organoid; *p < 0.05; ***p < 0.001; Student’s t-test). **(C)** Quantification of viral titers from CPO lysates (left) and culture supernatants (right) after S-CoV-2 treatment at 0, 24, and 72 hpi. Values represent mean ± SD (n = 2 biological replicates consisting of 4 organoids each; *p < 0.05; ***p < 0.001; Student’s t-test). FFU: focus forming units. **(D)** Representative confocal images of fluorescent immunohistology for DAPI, TUNEL, NP, and TTR in CPOs after S-CoV-2 or vehicle treatment at 24 and 72 hpi. Boxed regions highlight NP^+^TUNEL^+^ cells. Yellow arrows highlight NP^-^TUNEL^+^ cells near NP^+^ cells. Scale bar, 50 μm. **(E)** Quantification of percentages of TUNEL^+^NP^-^TTR^+^/NP^-^TTR^+^ cells (top) and TUNEL^+^NP^+^TTR^+^/NP^+^TTR^+^ cells (bottom) in CPOs after S-CoV-2 or vehicle treatment at 24 and 72 hpi. Values represent mean ± SEM with individual data points plotted (n = 3 organoids per hiPSC line with 3 images per organoid; ns: not significant; *p < 0.05; **p < 0.01; ***p < 0.001; Student’s t-test). Also see **Figure S3**.

Next, we examined the cellular consequence of SARS-CoV-2 infection of CPOs. We observed the presence of syncytia in SARS-CoV-2 infected cells, which significantly increased from an average of 4.6% at 24 hpi to 10.5% by 72 hpi (Figures S3D and S3E). The development of syncytia by cell-cell fusion mediated by interaction between SARS-CoV-2 spike protein and ACE2 expressed on adjacent cells has been reported with SARS-CoV-2 infection of several cell types and is a major mechanism by which the virus spreads to adjacent cells (Hoffmann et al., 2020a; Matsuyama et al., 2020; Ou et al., 2020). In particular, the SARS-CoV-2 spike protein appears to facilitate membrane fusion at a much higher rate than SARS-CoV-1 (Xia et al., 2020). By examining individual confocal Z-planes we could identify as many as 12 nuclei within a single infected cell at 72 hpi (Figure S3D). Interestingly, in addition to syncytia, we observed a significant increase in cell death in both uninfected and infected TTR^+^ choroid plexus cells in CPOs exposed to SARS-CoV-2 as measured by TUNEL immunolabeling (Figures 3D and 3E). Cell death increased from an average of 1.5% to 4.7% and 1.2% to 5.0% in uninfected and infected TTR^+^ choroid plexus cells from 24 to 72 hpi, respectively, across two hiPSC lines (Figure 3E). TUNEL^+^ uninfected cells tended to appear adjacent to SARS-CoV-2 infected cells, suggesting that infected cells may induce adjacent cells to die through an extrinsic mechanism (Figure 3D).

To confirm the susceptibility of choroid plexus epithelial cells to SARS-CoV-2 infection, we examined primary human choroid plexus epithelial cells (pHCPECs). pHCPECs exposed to SARS-CoV-2, but not vehicle control, for 12 hours showed many infected cells by immunolabeling with SARS-CoV-2 nucleoprotein at 72 and 120 hpi (Figure S3F). The number of infected cells increased with higher multiplicity of infection (MOI), with an average of 1.7% and 0.9% of cells infected at 72 and 120 hpi, respectively, using a MOI of 5 (Figure S3G).

Together, using CPOs as a model, our findings reveal SARS-CoV-2 tropism for choroid plexus epithelial cells that results in productive infection, increased cell syncytia that may promote viral spread through cell-cell fusion, and increased cell death among both infected and uninfected adjacent cells.

### Transcriptional Dysregulation of Choroid Plexus Organoids upon SARS-CoV-2 Infection

To gain additional insight into the functional consequence of SARS-CoV-2 infection in CPOs at the molecular level, we performed RNA-seq of SARS-CoV-2 and vehicle treated CPOs at 24 and 72 hpi. Principal component analysis showed clustering of biological replicates within different treatment groups (Figure S4A). After aligning reads to the SARS-CoV-2 genome, we detected high numbers of viral transcripts at 24 and 72 hpi, confirming that the virus was replicating within CPOs (Figure 4A). We also confirmed the expression of SARS-CoV-2 receptors ACE2 and NRP1 in CPOs at all time points (Figure 4B). Comparison of SARS-CoV-2 and vehicle treated CPOs revealed 1721 upregulated genes and 1487 downregulated genes at 72 hpi, indicating that SARS-CoV-2 infection leads to large scale transcriptional dysregulation (Figure S4B).

**Figure 4.**
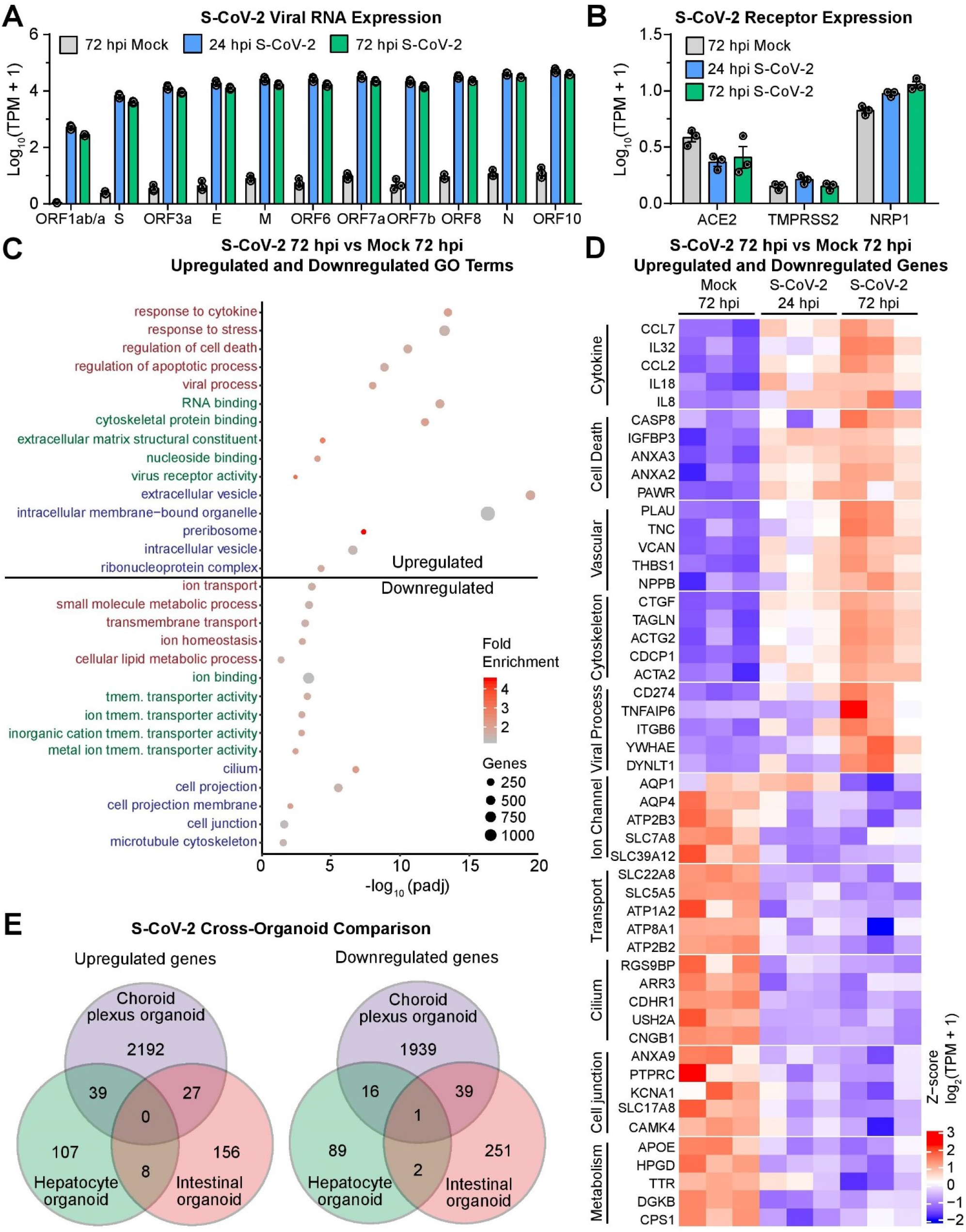
Transcriptional Dysregulation in Choroid Plexus Organoids upon SARS-CoV-2 Infection. **(A)** Quantification comparing Log_10_(TPM+1) of S-CoV-2 viral transcripts in CPOs after S-CoV-2 or vehicle treatment at 24 and 72 hpi by bulk RNA-seq. Values represent mean ± SEM with individual data points plotted (n = 3 biological replicates containing 3 organoids each). **(B)** Quantification comparing Log_10_(TPM+1) of known S-CoV-2 receptor transcripts in CPOs after S-CoV-2 or vehicle treatment at 24 and 72 hpi by RNA-seq. Values represent mean ± SEM with individual data points plotted (n = 3 biological replicates containing 3 organoids each). **(C)** Dot plot of selected enriched gene ontology (GO) terms for biological process (red), molecular function (green), and cellular component (blue) for upregulated and downregulated genes when comparing S-CoV-2 and vehicle treated CPOs at 72 hpi. **(D)** Heatmap of selected upregulated and downregulated genes when comparing S-CoV-2 and vehicle treated CPOs at 72 hpi. Genes related to different biological functions are grouped and labeled. Values are shown for each biological replicate as the row Z-score per gene of Log2(TPM+1)-transformed values. **(E)** Venn diagrams comparing the overlap of upregulated and downregulated genes following S-CoV-2 infection in CPOs, human hepatocyte organoids (Yang et al., 2020a), and human intestinal organoids (Lamers et al., 2020). Differentially expressed genes at both 24 and 72 hpi were combined for CPOs and differentially expressed genes for both expansion and differentiation intestinal organoid types were combined for intestinal organoids. Also see **Figure S4**.

More detailed gene ontology (GO) analysis of upregulated genes at 72 hpi showed enrichment for genes related to viral responses, RNA processing, response to cytokine, cytoskeletal rearrangement, and cell death (Figure 4C). Closer examination of upregulated genes revealed an increase in inflammatory cytokines CCL7, IL32, CCL2 (MCP1), IL18, and IL8 (Figure 4D). Notably, one case report examining inflammatory cytokines present in the serum and CSF of a patient with COVID-19 presenting with severe seizures identified CCL2 as the only cytokine enriched in the patient’s CSF compared to serum (Farhadian et al., 2020). CCL2 was one of the most highly upregulated genes after SARS-CoV-2 infection, suggesting that the choroid plexus may be the source of this cytokine in the CSF. IL32 plays important roles in both innate and adaptive immune responses by inducing TNF-alpha, which activates the signaling pathway of NF-kappa-B and p38 MAPK. This signaling pathway has been shown to be activated following SARS-CoV-2 infection in other cells (Bouhaddou et al., 2020). Upregulation of many vascular remodeling genes, such as NPPB, suggests that infected choroid plexus cells could signal the vasculature to promote leukocyte invasion (Shechter et al., 2013).

GO analysis of downregulated genes at 72 hpi, on the other hand, showed enrichment for genes related to ion transport, transmembrane transport, cilium, and cell junction (Figure 4C). Closer examination of downregulated genes revealed a decrease in the expression of many transporters and ion channels, such as AQP14 and SLC22A8, which are important for normal CSF secretory function (Figure 4D) (Brown et al., 2004; Hladky and Barrand, 2016), suggesting functional deficits of choroid plexus cells. The downregulation of many cell junction genes suggests a remodeling or break-down of normal CSF-blood barrier function, which also occurs during brain inflammation (Engelhardt and Tietz, 2015). Additionally, the dramatic decrease in TTR production may result in decreased thyroxine transport to the CSF, which has been shown to contribute to neuropsychiatric symptoms and “brain fog” or difficulty in concentrating in patients (Samuels, 2014).

Transcriptional dysregulation induced by SARS-CoV-2 in CPOs at 24 hpi showed similar gene expression changes as at 72 hpi, although less severe in most cases (Figures 4D and S4C-D). Comparison of the list of dysregulated genes at 24 and 72 hpi revealed a substantial overlap of 42% and 32% of upregulated and downregulated genes, respectively (Figure S4E).

To investigate whether organoids from different tissues exhibit similar transcriptional responses upon SARS-CoV-2 infection, we compared SARS-CoV-2 induced transcriptional changes among CPOs and published datasets from hepatocyte organoids (Yang et al., 2020a) and intestinal organoids (Lamers et al., 2020). Surprisingly, we found little overlap among upregulated or downregulated genes in the three different organoid types, suggesting a predominantly cell type-specific response to SARS-CoV-2 infection (Figure 4E). This finding supports the use of a diverse array of human model systems to investigate effects of SARS-CoV-2 infection. Additionally, we found many more dysregulated genes in CPOs compared to the other two organoid types, suggesting that SARS-CoV-2 may exhibit a greater impact on choroid plexus cells (Figure 4E).

Together, these transcriptome analyses suggest that SARS-CoV-2 infection of choroid plexus cells in CPOs leads to viral RNA production and inflammatory cellular responses with a concomitant compromise of normal secretory functions and increased cell death. Furthermore, responses to SARS-CoV-2 infection are largely tissue organoid specific, at least at the transcriptional level.

## DISCUSSION

Using a platform of hiPSC-derived monolayer neurons, microglia, astrocytes, and region-specific brain organoids, we showed modest tropism of SARS-CoV-2 for neurons and astrocytes with a particularly high rate of infection of choroid plexus epithelial cells. Using an optimized protocol for generating choroid plexus organoids (CPOs) from hiPSCs that resemble the morphology, marker expression, and transcriptome of human choroid plexus, we revealed productive infection of SARS-CoV-2 in choroid plexus epithelial cells, which we confirmed using primary human choroid plexus epithelial cells. Analysis of the consequence of SARS-CoV-2 infection of CPOs showed an increase in both cell autonomous and non-cell autonomous cell death and transcriptional dysregulation related to increased inflammation and altered secretory function. Our study provides an organoid platform to investigate the cellular susceptibility, pathogenesis, and treatment strategies of SARS-CoV-2 infection of brain cells and further implicates the choroid plexus as a potentially important site for pathology.

### Modeling SARS-CoV-2 Infection with Brain Organoids

Although COVID-19 primarily manifests as respiratory distress resulting from inflammatory damage to the lung epithelium, there is mounting clinical evidence implicating other organs, such as the kidney, intestine, and brain, in disease pathogenesis (Renu et al., 2020). Because SARS-CoV-2 can affect multiple organs, the currently widely used cellular model systems, such as Vero E6 cells or human cancer cell lines, do not adequately recapitulate the cellular diversity and breadth of the disease. Although mice engineered to express human ACE2 have provided an *in vivo* model to study SARS-CoV-2, the system relies on human ACE2 overexpression and does not fully recapitulate symptoms of COVID-19 in humans. Recently, organoids have emerged as a model to study human cells and organogenesis *in vitro* (Clevers, 2016). Organoids for the gut, kidney, endothelium, liver, and pancreas have been used to better understand SARS-CoV-2 viral tropism and pathology (Lamers et al., 2020; Monteil et al., 2020; Yang et al., 2020a; Zhou et al., 2020). These studies support cell-type specific susceptibility to SARS-CoV-2 infection. Our finding that dysregulated gene expression varies widely among hepatocyte, intestinal, and choroid plexus organoids infected with SARS-CoV-2 suggests unique responses in different cell types and highlights the need for diverse cellular model systems when studying the disease. Brain organoid models for SARS-CoV-2 infection are particularly useful since it is not feasible to take brain biopsies from patients with COVID-19. The load of SARS-CoV-2 that may enter the brain is likely to be inconsistent throughout the course of illness and may be difficult to accurately assess in post-mortem samples. For example, viral particles or RNA has been detected in the CSF or brain of some but not the majority of patients with neurological symptoms (Helms et al., 2020; Moriguchi et al., 2020; Puelles et al., 2020; Solomon et al., 2020; Virani et al., 2020), although the amount of viral particles in the CSF may be insufficient for detection (Ye et al., 2020). Brain organoids allow to observe the acute and long-term impact of SARS-CoV-2 infection in real time, with the ability to introduce perturbations, such as therapeutic agents, to assess cellular responses. The use of region-specific brain organoids, including CPOs, expands the organoid toolset for investigating SARS-CoV-2 infection, and provides a platform for testing new therapeutics.

Building upon previous findings (Sakaguchi et al., 2015; Watanabe et al., 2012), we optimized a CPO model that is simple, robust and reproducible with the initial goal to study SARS-CoV-2 infection. These CPOs express high levels of SARS-CoV-2 receptors and exhibit productive infection of SARS-CoV-2 and pathogenesis at cellular and molecular levels. CPOs may provide a better model system than primary human choroid plexus epithelial cells, which are not widely available and often lose some characteristics upon multiple rounds of expansion in culture and showed a lower infection rate than our CPOs. Unlike a recently developed protocol that utilizes cultures with Matrigel extracellular matrix (Pellegrini et al., 2020), our CPOs do not obviously polarize or develop fluid-filled, cystic structures, but do offer the benefit of not relying on Matrigel, which has substantial batch-to-batch variation and can lead to increased inter-organoid variability. Our protocol uses feeder-free hiPSCs to improve organoid reproducibility, and CPOs are cultured on an orbital shaker to enhance oxygen and nutrient diffusion throughout organoids. With these improvements, CPOs reproducibly generate projections of cuboidal cells and exhibit uniform expression of choroid plexus markers OTX2, AQP1, and TTR in most of the cells within organoids. They also exhibit a transcriptome similar to adult human choroid plexus tissue. Importantly, these CPOs express many transporters that are essential for CSF secretion. These CPOs provide a platform for future investigations of cell type-specific pathogenesis, mechanisms and treatment of SARS-CoV-2 infection. Our CPO platform can also be used to model human choroid plexus development and associated brain disorders in the future (Lun et al., 2015).

### Neurotropism of SARS-CoV-2 and Choroid Plexus as a Potential Site for Viral Entry and Spread in the CNS

ACE2 has been identified as a key cell entry receptor for SARS-CoV-2 (Hoffmann et al., 2020b). Although moderate ACE2 expression has been detected in various brain regions, particularly high levels have been reported in the choroid plexus in humans and mice (Chen et al., 2020). The high levels of ACE2 expression in CPOs may explain their higher susceptibility to SARS-CoV-2 infection compared to other brain organoids. Moreover, the productive infection and high incidence of syncytia in infected choroid plexus cells may contribute to the viral spread. On the other hand, despite relatively low ACE2 expression, we did detect sparse infection of neurons by SARS-CoV-2. Recent studies point to the potential for other proteins that are expressed more highly in neural cells, such as NRP1, to serve as receptors for SARS-CoV-2. We did not observe obvious increase in the number of infected neurons over time, suggesting that the virus does not spread efficiently from neuron-to-neuron.

Central nervous system disorders, including seizures and encephalopathy, have been reported with SARS-CoV-1 with detection of infectious virus and RNA in the CSF (Hung et al., 2003; Lau et al., 2004; Xu et al., 2005). Similar findings are present in patients with COVID-19 (Helms et al., 2020; Moriguchi et al., 2020; Puelles et al., 2020; Solomon et al., 2020; Virani et al., 2020), but how SARS-CoV-2 may enter the brain is currently unknown. The main hypotheses are either neuron-to-neuron spread via bipolar cells located in the olfactory epithelium that extend axons and dendrites to the olfactory bulb, or a hematogenous route across the blood-CSF-barrier (Desforges et al., 2014). Neuron to neuron propagation has been described for other coronaviruses (Dube et al., 2018; Netland et al., 2008), but it was not obvious in our study. Our finding that the choroid plexus is particularly susceptible to productive infection with SARS-CoV-2, raises the possibility that choroid plexus epithelial cells could be infected from virus circulating in closely associated capillaries. SARS-CoV-2 may replicate within choroid plexus cells and shed viral copies into the CSF. Indeed, we detected increased viral load in culture supernatants overtime. Alternatively, virus may enter the CSF via the cribiform plate in nasal passages and replicate in choroid plexus cells, increasing viral availability in the central nervous system. Viral propagation through the CSF may explain cases of spinal cord pathology in some patients with COVID-19 (Giorgianni et al., 2020; Valiuddin et al., 2020). More detailed analyses of post-mortem brain samples from patients with COVID-19 and animal models of SARS-CoV-2 infection are necessary to better understand potential routes of viral entry into the brain.

### Potential Role of the Choroid Plexus in SARS-CoV-2 Pathogenesis

The transcriptional dysregulation in CPOs after SARS-CoV-2 infection indicates an inflammatory response and perturbed choroid plexus cellular function. Many pro-inflammatory cytokines were upregulated, including CCL7, IL32, CCL2, IL18, and IL8, which are important for recruiting immune cells to sites of infection. Upregulation of vascular and extracellular matrix remodeling genes also suggests that during infection, choroid plexus cells can signal the adjacent vasculature to recruit inflammatory cells. Downregulation of many genes important for CSF secretion, including AQP1, and many ion transporters, such as ATP1A2, may lead to altered CSF production and composition. Mutations in many of these dysregulated transporter genes have been implicated in neurological complications in humans, such as migraine and encephalopathy (Murphy et al., 2018). Increased expression of cell death genes supports our finding of increased numbers of TUNEL^+^ cells after SARS-CoV-2 infection. Increased cell death and downregulation of cell junction genes, such as CLDN2, suggest damage to tight junctions between choroid plexus epithelial cells, which may disturb the blood-CSF-barrier, allowing pathogens and regulated solutes to pass more freely (Ghersi-Egea et al., 2018). Downregulation of many genes related to the cilium suggests an impaired cytoskeleton and dysfunctional sensing of the extracellular environment. There was also a significant downregulation of TTR, which is necessary for carrying thyroid hormone from the blood into the CSF, and lowered TTR production by the choroid plexus has been associated with neuropsychiatric disorders, such as schizophrenia and depression (Huang et al., 2006; Sullivan et al., 1999). Furthermore, low CSF thyroid hormone has been associated with many neurological symptoms, including slowing of thought and speech, decreased attentiveness, and reduced cognition, which have all been reported by patients with COVID-19 (Ellul et al., 2020). Our study implicates the choroid plexus epithelium as a potential target of SARS-CoV-2 infection and a contributor to COVID-19 disease pathogenesis that warrants further investigation.

### Limitations of *in vitro* Neural Cultures to Model Infection

Our study demonstrates clear susceptibility of hiPSC-derived neurons, astrocytes, and choroid plexus cells, as well as primary astrocytes and choroid plexus epithelial cells, to SARS-CoV-2 infection *in vitro*, however, a thorough analysis of brain samples from patients with COVID-19 is necessary to confirm neural susceptibility *in vivo*. Our methodology has a number of constrains that limit result interpretation. First, since brain organoids more closely resemble the developing fetal brain than the mature adult brain, there may exist important differences in SARS-CoV-2 susceptibility between immature and mature cells. There has been recent evidence of vertical transmission of SARS-CoV-2 from mother to fetus (Dong et al., 2020; Vivanti et al., 2020; Zeng et al., 2020), but impact on the fetal brain remains unclear. Brain organoids may serve as a useful model system to explore the potential effects of fetal SARS-CoV-2 exposure on brain development. Second, direct exposure of brain organoid cultures to SARS-CoV-2 in the culture medium may not accurately recapitulate the physiological amount and duration of viral exposure in humans. Brain organoid models also lack an intact blood-brain-barrier or blood-CSF-barrier which may modulate SARS-CoV-2 access to the brain. Third, our brain organoids lack additional cell types, such as oligodendrocytes, stromal cells, immune cells, and endothelial cells, which may also modulate susceptibility to infection and contribute to disease pathogenesis *in vivo*. Immune cells, such as T cells and monocytes, have been shown to mediate the host response to SARS-CoV-2 infection, including tissue-specific inflammation (Tay et al., 2020). Endothelial cells are major targets of SARS-CoV-2 and may be the primary cause of SARS-CoV-2-related effects in the brain with neurological symptoms being secondary to vascular changes and hypoxia (Ellul et al., 2020). Extending studies of SARS-CoV-2 susceptibility to a more diverse selection of brain cells that includes neurons of the peripheral nervous system is an important future direction.

It is currently unclear if the neurological symptoms present in patients with COVID-19 are a direct result of neural infection or secondary to endothelial cell damage and hypoxia, circulating pro-inflammatory cytokines, or ambulatory treatment. Further studies in humans will be necessary to better correlate the onset and severity of neurological symptoms with neural cell infection. Our *in vitro* studies provide useful information on which specific cell types to focus on for future human studies and offers a simple, accessible and tractable human cell platform to investigate the cellular susceptibility, disease mechanisms, and treatment strategies for SARS-CoV-2 infection of the human brain.

## ACKNOWLEDGEMENTS

We thank members of Ming and Song laboratories for comments and suggestions, B. Temsamrit and E. LaNoce for technical support, and J. Schnoll for lab coordination. This work was supported by grants from National Institutes of Health (U19AI131130 and R35NS097370 to G-l.M., U19MH106434 to A.G.B, H.S. and G-l.M., and R35NS116843 to H.S.), the Perelman Professor of Neuroscience Endowment Fund from University of Pennsylvania Perelman School of Medicine (to H.S. and G-l.M.), and by the Intramural Research Programs of the National Center for Advancing Translational Sciences (to W.Z.).

## AUTHOR CONTRIBUTIONS

F.J. contributed to histological analysis of various brain organoids and development of the choroid plexus organoid model. S.R.P. and F.Z. contributed to RNA-seq data analysis. W-K.H. and S.Z.H.W. performed hypothalamic organoid analysis. F.Z. and D.Y.Z. performed RNA-seq. H.Z. contributed to monolayer cell culture, immunocytochemistry and analyses. B.C. contributed to preparation of viral infected samples and viral titer analysis. C.Z.C., M.X., M.P., W.Z. contributed to midbrain organoid generation. J.C.T. performed viral infection. W.Z., A.G.B., H.S., J.C.T., G-l.M. conceived and guided the project. F.J., H.S. and G-l.M. wrote the manuscript with inputs from all authors.

---

**Figure S1.**
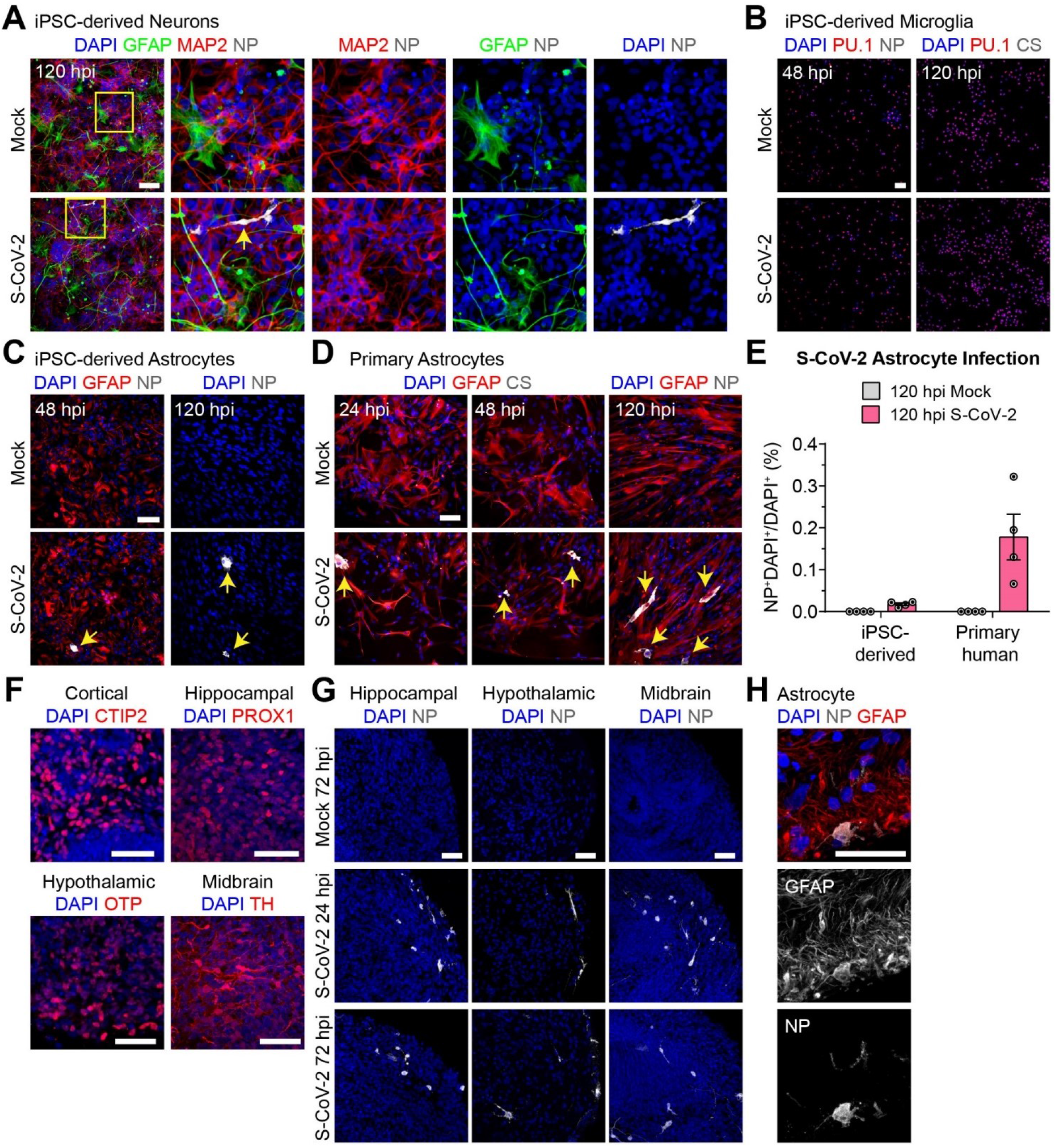
SARS-CoV-2 Neurotropism in hiPSC-derived Monolayer Neural Cultures and Brain Organoids, related to Figure 1. **(A)** Representative confocal images of fluorescent immunocytochemistry for DAPI, SARS-CoV-2 (S-CoV-2) nucleoprotein (NP), GFAP, and MAP2 in hiPSC-derived cortical neurons plated on hiPSC-derived astrocytes after S-CoV-2 or vehicle treatment at 120 hpi. Arrow points to an infected MAP2+ neuron. Scale bar, 50 μm. **(B)** Representative confocal images of fluorescent immunocytochemistry for DAPI, NP or convalescent serum from a patient with COVID-19 (CS), and PU.1 in hiPSC-derived microglia after S-CoV-2 or vehicle treatment at 48 and 120 hpi. Scale bar, 50 μm. Note a lack of infected cells at both time points. **(C)** Representative confocal images of fluorescent immunocytochemistry for DAPI, NP, and GFAP in hiPSC-derived astrocytes after S-CoV-2 ore vehicle treatment at 48 and 120 hpi. Scale bar, 50 μm. Arrows point to infected cells. **(D)** Representative confocal images of fluorescent immunocytochemistry for DAPI, NP or patient convalescent serum (CS), and GFAP in human primary astrocytes after S-CoV-2 or vehicle treatment at 24, 48, and 120 hpi. Scale bar, 50 μm. Arrows point to infected cells. **(E)** Quantification of percentages of NP+DAPI+/DAPI+ cells in mock and S-CoV-2 treated hiPSC-derived astrocyte and human primary astrocyte cultures at 120 hpi. Values represent mean ± SEM with individual data points plotted (n = 4 independent wells per condition). **(F)** Representative confocal images of fluorescent immunohistology for DAPI and CTIP2, PROX1, OTP, and TH in cortical, hippocampal, hypothalamic, and midbrain organoids, respectively. Scale bars, 50 μm. **(G)** Representative confocal images of fluorescent immunohistology for DAPI and NP in hippocampal, hypothalamic, and midbrain organoids derived from a second independent hiPSC line after S-CoV-2 ore vehicle treatment at 24 and 72 hpi. Scale bars, 50 μm. **(H)** Representative confocal images of fluorescent immunohistology for DAPI, NP, and GFAP in a hypothalamic organoid after S-CoV-2 treatment at 72 hpi. Scale bar, 50 μm.

**Figure S2.**
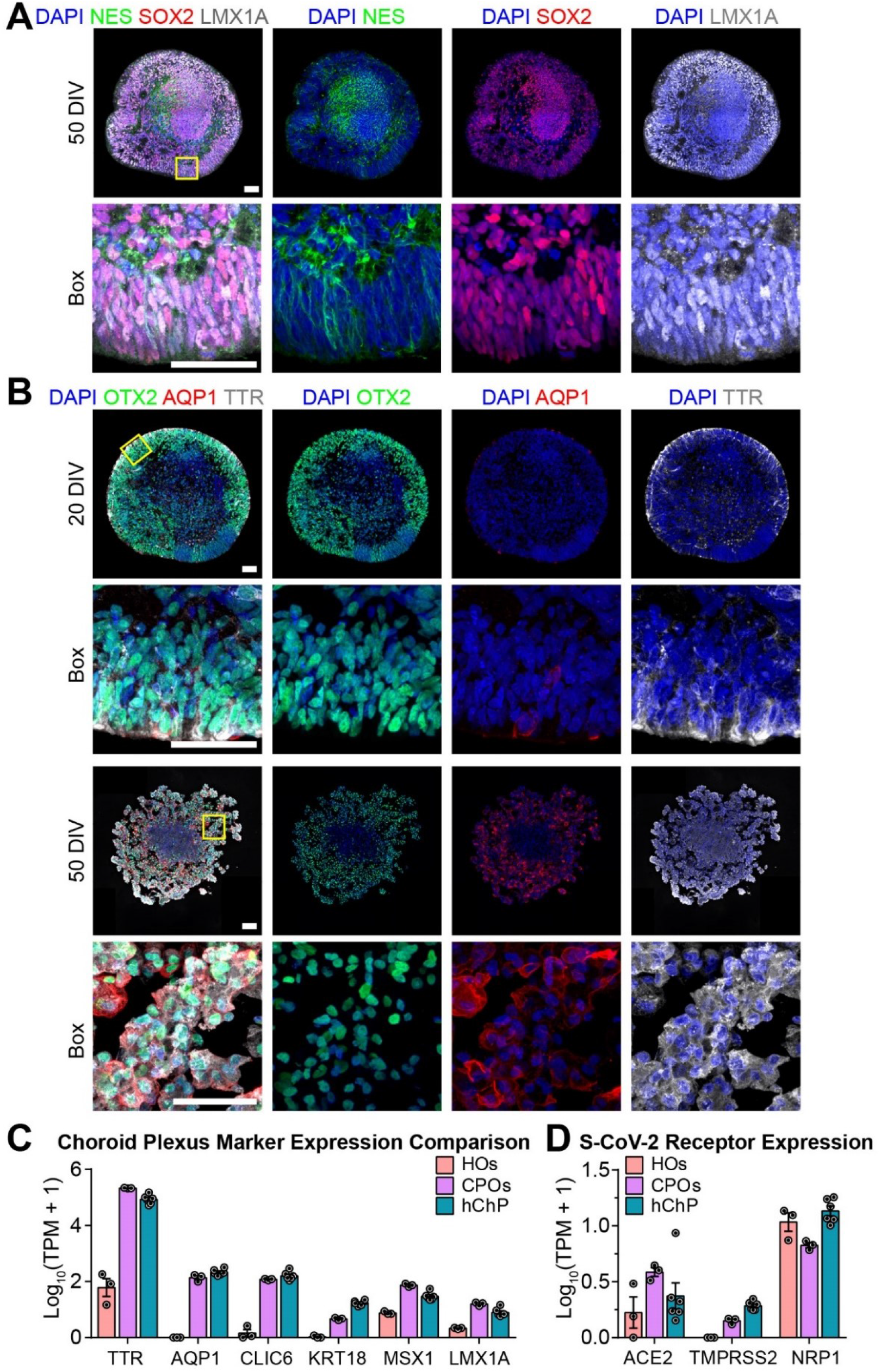
Additional Characterization of hiPSC-derived Choroid Plexus Organoids, related to Figure 2. **(A)** Representative confocal images of fluorescent immunohistology for DAPI, LMX1A, NES, and SOX2 in CPOs at 20 DIV for a second hiPSC line. Scale bars, 50 μm. **(B)** Representative confocal images of fluorescent immunohistology for DAPI, OTX2, AQP1, and TTR in CPOs at 20 and 50 DIV for a second hiPSC line. Scale bars, 50 μm. **(C)** Quantification of the Log_10_(TPM+1) expression of choroid plexus marker genes in 45 DIV hippocampal organoids (HOs), 50 DIV CPOs, and adult human choroid plexus tissue (hChP) (Rodriguez-Lorenzo et al., 2020) by bulk RNA-seq. Values represent mean ± SEM with individual data points plotted (n = 3 biological replicates for organoids and n = 6 biological replicates for the human choroid plexus tissue). **(D)** Quantification of the Log_10_(TPM+1) expression of known S-CoV-2 receptors in 45 DIV HOs, 50 DIV CPOs, and adult human choroid plexus tissue (hChP) by bulk RNA-seq. Values represent mean ± SEM with individual data points plotted (n = 3 biological replicates for organoids and n = 6 biological replicates for human choroid plexus).

**Figure S3.**
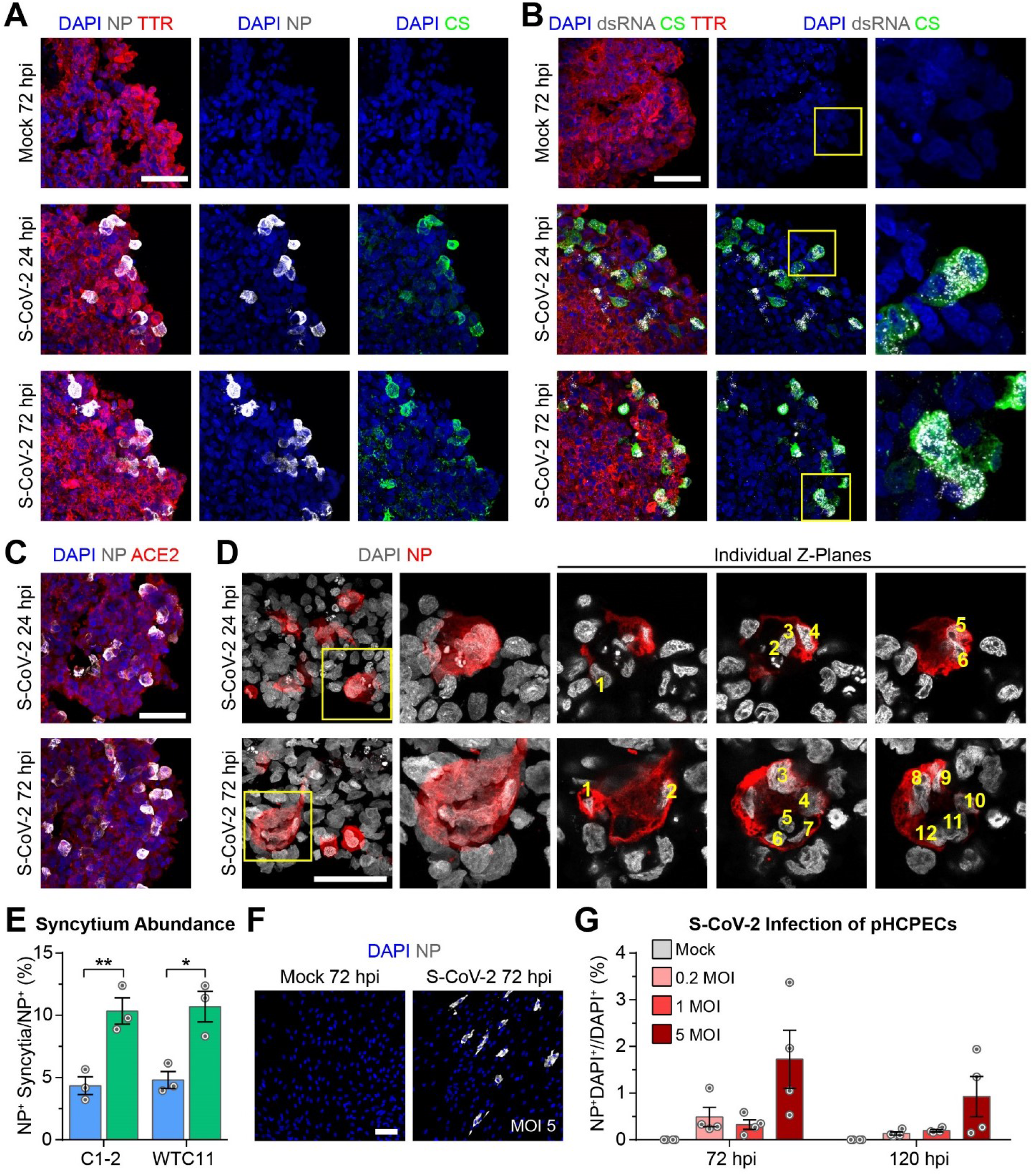
Additional Characterization of SARS-CoV-2 Infected Choroid Plexus Organoids and Infection of Primary Human Choroid Plexus Epithelial Cells, related to Figure 3. **(A)** Representative confocal images of fluorescent immunohistology for DAPI, SARS-CoV-2 nucleoprotein (NP), patient convalescent serum (CS), and TTR in CPOs after S-CoV-2 or vehicle treatment at 24 and 72 hpi in a second hiPSC line. Scale bar, 50 μm. **(B)** Representative confocal images of fluorescent immunohistology for DAPI, double-stranded RNA (dsRNA), patient convalescent serum (CS), and TTR in CPOs after S-CoV-2 or vehicle treatment at 24 and 72 hpi. Boxed regions highlight NP^+^ cells containing many dsRNA puncta. Scale bar, 50 μm. **(C)** Representative confocal images of fluorescent immunohistology for DAPI, NP, and ACE2 in CPOs after S-CoV-2 treatment at 24 and 72 hpi. Scale bar, 50 μm. **(D)** Representative confocal images of fluorescent immunohistology for DAPI and NP in CPOs after S-CoV-2 treatment at 24 and 72 hpi. Boxed regions highlight NP^+^ syncytia with individual Z-planes separated to highlight multiple nuclei counted within each syncytium. Scale bar, 50 μm. **(E)** Quantification of the percentages of NP^+^ syncytia/Total NP^+^ cells in CPOs after S-CoV-2 treatment at 24 and 72 hpi. Values represent mean ± SEM with individual data points plotted (n = 3 organoids per hiPSC line with 3 images per organoid; *p < 0.05; **p < 0.01; Student’s t-test). **(F)** Representative confocal images of fluorescent immunohistology for DAPI and NP in primary human choroid plexus epithelial cell (pHCPEC) cultures after S-CoV-2 or vehicle treatment at 72 hpi. Scale bar, 50 μm. **(G)** Quantification of the percentages of NP^+^DAPI^+^/DAPI^+^ cells in pHCPEC cultures after S-CoV-2 or vehicle treatment using different MOIs at 72 and 120 hpi. Values represent mean ± SEM with individual data points plotted (n = 4 individual wells).

**Figure S4.**
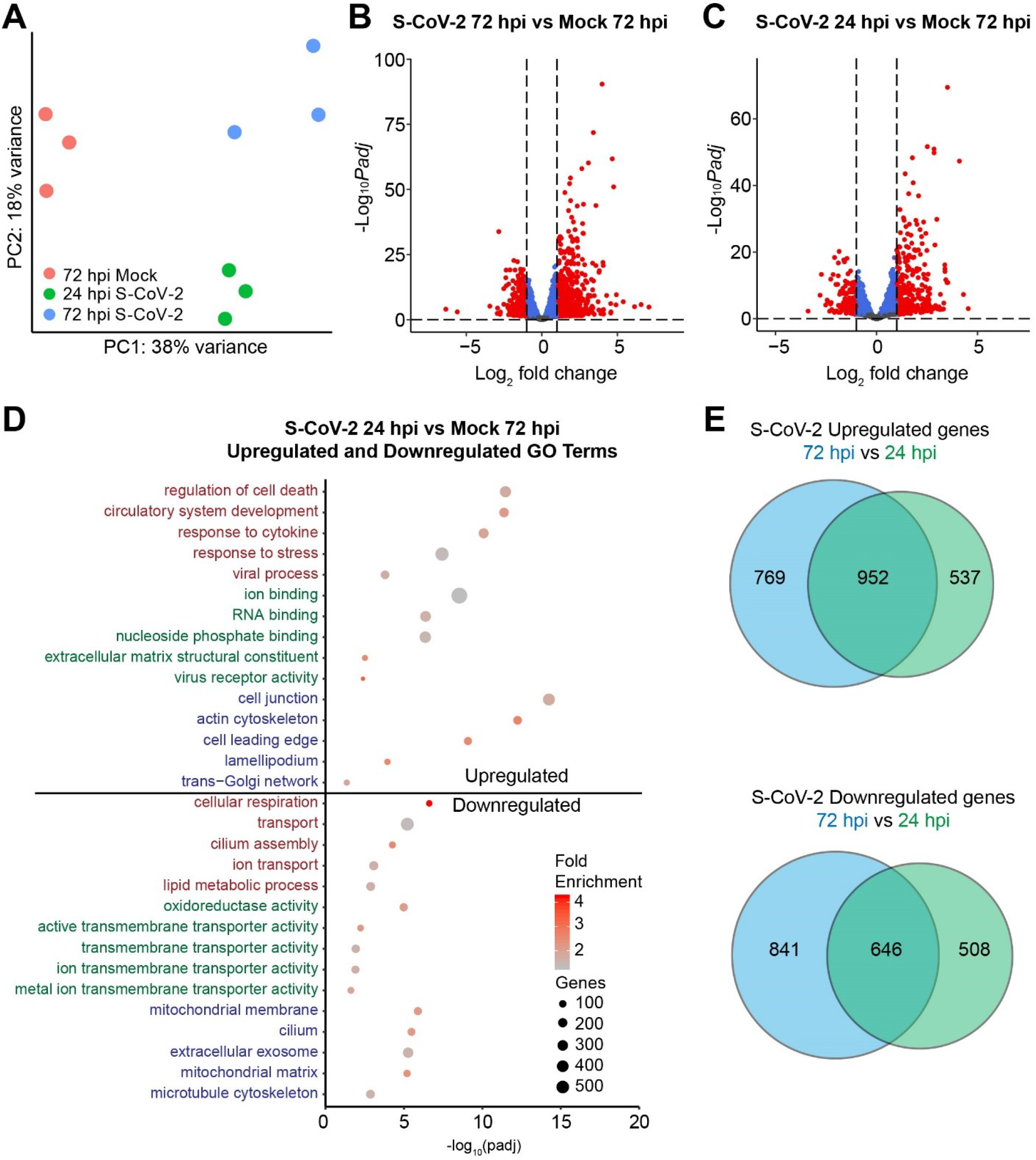
Additional analysis of transcriptional dysregulation in SARS-CoV-2 Infected Choroid Plexus Organoids, related to Figure 4. **(A)** Principal component analysis (PCA) plot comparing the bulk RNA transcriptomes of CPOs after S-CoV-2 or vehicle treatment at 24 and 72 hpi. Three biological replicates are shown for each condition. **(B)** Volcano plot showing the differentially expressed genes comparing S-CoV-2-treated and vehicle-treated CPOs at 72 hpi. The gene expression data for 3 biological replicates for each condition were averaged before comparison. Red dots represent genes with log_2_(fold change)> |1| and *Padj* < 0.05. **(C)** Volcano plot showing the differentially expressed genes comparing S-CoV-2-treated CPOs at 24 hpi and vehicle-treated CPOs at 72 hpi. The gene expression data for 3 biological replicates for each condition were averaged before comparison. Red dots represent genes with log_2_(fold change)> |1| and *Padj* < 0.05. **(D)** Dot plot of selected enriched gene ontology (GO) terms for biological process (red), molecular function (green), and cellular component (blue) for upregulated and downregulated genes when comparing S-CoV-2-treated CPOs at 24 hpi and vehicle-treated CPOs at 72 hpi. **(E)** Venn diagrams comparing the overlap of upregulated (top) and downregulated (bottom) genes in S-CoV-2-treated CPOs at 24 and 72 hpi compared to the vehicle control.

## METHODS

### Human Induced Pluripotent Stem Cells

Human iPSC lines used in the current study were derived from healthy donors and were either obtained from commercial sources or previously generated and fully characterized (Chiang et al., 2011; Wen et al., 2014; Yoon et al., 2014). C1-2 hiPSCs were generated from fibroblasts obtained from ATCC (CRL-2097). C3-1 hiPSCs were generated from fibroblasts obtained from an individual in a previously characterized American family (Sachs et al., 2005). WTC11 hiPSC were obtained from Coriell (GM25256). HT268A hiPSCs were generated from dermal fibroblasts obtained from Coriell (GM05659) (Vu et al., 2018), and PCS201012 hiPSCs were generated from dermal fibroblasts obtained from ATCC (PCS-201-012) (Beers et al., 2015). Generation of iPSC lines followed institutional IRB and ISCRO guidelines and was approved by Johns Hopkins University School of Medicine. Karyotyping analysis by standard G-banding technique was carried out by the Cytogenetics Core Facility at the Johns Hopkins Hospital or Cell Line Genetics. Results were interpreted by clinical laboratory specialists of the Cytogenetics Core or Cell Line Genetics. All hiPSC lines were confirmed free of Mycoplasma, Acholeplasma, and Spiroplasma with a detection limit of 10 CFU/ml by targeted PCR (Biological Industries).

### Monolayer Neural Cell Cultures

hiPSC-derived cortical astrocytes (BrainXell, Inc.) and microglia (BrainXell, Inc.) were cultured according to the supplier’s instructions and plated directly out of thaw onto 96 well imaging plates coated with Poly-L-ornithine at 2 μg/cm^2^ and laminin from Engelbreth-Holm-Swarm murine sarcoma basement membrane at 5 μg/mL at a density of 15,000 cells per well. hiPSC-derived cortical glutamatergic neurons, cortical GABAergic neurons, and cortical astrocytes (BrainXell, Inc.) were plated according to the supplier’s instructions, mixed together in a ratio of 10,000 GABAergic neurons:10,000 Glutamatergic neurons:5000 astrocytes directly out of thaw onto 96 well imaging plates coated with Poly-L-ornithine (Sigma-Aldrich) at 2 μg/cm^2^ and laminin from Engelbreth-Holm-Swarm murine sarcoma basement membrane (Sigma-Aldrich) at 5 μg/mL. The day after plating the media was removed, and replaced with BrainPhys medium (StemCell Technologies) with 1% N2 (ThermoFisher Scientific), 2% B27 (ThermoFisher Scientific), 1% Pen-strep (ThermoFisher Scientific), and supplemented with recombinant Brain-derived Neurotrophic Factor (BDNF, 20 ng/ml; Peprotech), recombinant Glia-derived Neurotrophic Factor (GDNF, 20 ng/ml; Peprotech), ascorbic acid (200 nM; Sigma), dibutyryl cyclic AMP (1 mM; Sigma) and laminin (1 μg/mL; Sigma-Aldrich) (Bardy et al., 2015). Primary human astrocytes (ScienCell Research Laboratories) were cultured and expanded for two passages, according to the supplier’s instructions. After the second expansion, astrocytes were cryopreserved at 250,000 cells per vial. Banked astrocytes were thawed directly onto 96 well imaging plates (Greiner Bio-one) coated with Poly-L-ornithine (Sigma-Aldrich) at 2 μg/cm^2^ and laminin from Engelbreth-Holm-Swarm murine sarcoma basement membrane (Sigma-Aldrich) at 5 μg/mL at a density of 16,000 cells per well for shorter infection times (48 hours or less) and 8000 cell per well for 120 hour infections. Primary human choroid plexus epithelial cells (ScienCell Research Laboratories) were cultured according to the supplier’s instructions. Cells were either passaged once or plated directly out of thaw onto 96-well imaging plates coated with Poly-L-ornithine at 2 μg/cm^2^ at a density of 4000 cells per well.

### Human Induced Pluripotent Stem Cell Culture

All studies involving hiPSCs were performed under approved protocols of the University of Pennsylvania and NIH. All hiPSC lines were confirmed to have a normal karyotype and were confirmed free of Mycoplasma, Acholeplasma, and Spiroplasma with a detection limit of 10 CFU/mL by targeted PCR. For cortical, hippocampal, hypothalamic, and midbrain organoid generation, hiPSCs were cultured on mouse embryonic fibroblast feeder (MEF) cells in stem cell medium consisting of DMEM:F12 supplemented with 20% KnockOut Serum Replacement, 1X MEM-NEAAs, 1X GlutaMAX, 1X Penicillin-Streptomycin, 1X 2-mercaptoethanol, and 10 ng/mL bFGF in a 5% CO_2_, 37°C, 90% relative humidity incubator as previously described (Qian et al., 2018). Culture medium was replaced every day. hiPSCs were passaged every week onto a new tissue-treated culture plate coated with 0.1% gelatin for 2 hours and pre-seeded with γ-irradiated CF1 MEF cells one day in advance. Colonies were detached by washing with DPBS and treating with 1 mg/mL Collagenase Type IV for 30-60 minutes. Detached colonies were washed 3 times with 5 mL DMEM:F12 and dissociated into small clusters by trituration with a P1000 pipette. For choroid plexus organoid generation, hiPSCs were maintained in feeder-free conditions on plates pre-coated with hESC-qualified Matrigel (Corning) using mTeSR Plus medium (Stemcell Technologies). Culture medium was replaced every other day and hiPSCs were passaged every week by washing with DPBS and incubating with ReLeSR reagent (Stemcell Technologies) for 5 minutes. Detached hiPSCs were broken into smaller clusters by gentle trituration using a 5 mL pipette and seeded onto fresh coated plates.

### Brain Organoid Cultures

Generation of cortical organoids from C3-1 hiPSCs was performed as previously described (Qian et al., 2018; Qian et al., 2016). Briefly, hiPSC colonies were detached with Collagenase Type IV, washed with DMEM:F12, and transferred to an ultra-low attachment 6-well plate (Corning Costar) in neural induction medium containing DMEM:F12 supplemented with 20% KnockOut Serum Replacement, 1X MEM-NEAAs, 1X GlutaMax, 1X Penicillin-Streptomycin, 1X 2-mercaptoethanol, 2 μM Dorsomorphin and 2 μM A83-01. On Day 5 and Day 6, half of the medium was replaced with forebrain induction medium consisting of DMEM:F12 supplemented with 1X N2 supplement, 1X Penicillin/Streptomycin, 1X Non-essential Amino Acids, 1X GlutaMax, 1 μM CHIR99021, and 1 μM SB-431542. On Day 7, organoids were embedded in Matrigel and cultured in forebrain induction medium for 7 more days. On Day 14, embedded organoids were mechanically dissociated from Matrigel and transferred to a 6 well-plate on a CO_2_ resistant orbital shaker (ThermoFisher) and grown in differentiation medium consisting of DMEM:F12 supplemented with 1X N2 supplement, 1X B27 supplement, 1X Penicillin/Streptomycin, 1X 2-Mercaptoethanol, 1X Non-essential Amino Acids, and 2.5 g/mL human insulin. From Day 35 until the point of sampling, extracellular matrix (ECM) proteins were supplemented in differentiation medium by dissolving Matrigel at 1% (v/v). To generate midbrain organoids from HT268A and PCS201012 hiPSCs, hiPSC colonies were transferred to an Ultra-Low attachment 6-well plate containing Midbrain Induction Medium 1, consisting of DMEM:F12, 20% Knockout Serum Replacer, 1X Glutamax, 1X 2-Mercaptoenthanol, 1x Pen/Strep, 100 nM LDN-193189, 10 μM SB-431542, 2 μM Purmorphamine, 100 ng/mL SHH, 100 ng/mL FGF-8. On day 5, cultures were switched to Midbrain Induction Medium 2, consisting of DMEM:F12, 1X N2 Supplement, 1X Glutamax, 1x Pen/Strep, 100 nM LDN-193189, 3 μM CHIR-99021, 2 μM Purmorphamine, 100 ng/ml SHH, 100 ng/ml FGF-8. On day 7, EBs with smooth and round edges were selected, switched to Midbrain Induction Medium 3, consisting of DMEM:F12, 1X N2 Supplement, 1X Glutamax, 1x Pen/Strep, 100 nM LDN-193189, 3 μM CHIR-99021, and transferred to a shaking platform. On day 14, cultures were switched to Differentiation Medium, consisting of Neurobasal Medium, 1X B27 Supplement, 1X Glutamax, 1X 2-Mercaptoenthanol, 20 ng/mL BDNF, 20 ng/mL GDNF, 0.2 mM Ascorbic Acid, 1 ng/mL TGFβ, and 0.5 mM c-AMP. For generation of hippocampal organoids from C1-2 and WTC11 hiPSC lines, cortical induction medium was replaced with medium containing 3 μM CHIR-99021 and 20 ng/mL recombinant human BMP-7. For generation of hypothalamic organoids from C1-2 and C3-1 hiPSC lines, hiPSC colonies were processed similarly to those for cortical organoids followed by exposure to Shh signaling and Wnt-inhibition in induction medium with 50 ng/mL recombinant Sonic Hedgehog, 1 μM Purmorphamine, 10 μM IWR1-endo and 1 μM SAG, and then transferred to differentiation medium for further culture. Organoids were maintained with media changes every other day.

### Choroid Plexus Organoid Culture

At 70% confluence, undifferentiated hiPSCs (C1-2 and WTC11 lines) grown in feeder-free conditions were detached using ReLeSR reagent and gently triturated into small clusters using a 5 mL pipette and pipette aid. hiPSC clusters were centrifuged at 200 g for 3 minutes and the supernatant was aspirated and replaced with mTeSR Plus medium supplemented with 10 μM Y-27632. Approximately 5,000 hiPSCs were aggregated into each embryoid body by following the instructions using the Aggrewell 800 plate and cultured overnight in mTeSR Plus medium supplemented with 10 μM Y-27632. The next day (Day 1), embryoid bodies were gently removed from the Aggrewell plate using a P1000 pipettor with a wide-bore tip, washed three times with DMEM:F12, and transferred to neural induction medium containing DMEM:F12 supplemented with 20% KnockOut Serum Replacement, 1X MEM-NEAAs, 1X GlutaMax, 1X Penicillin-Streptomycin, 1X 2-mercaptoethanol, 0.5 μM LDN-193189, 5 μM SB-431542, 1 μM IWP-2, and 10 μM Y-27632 in ultra-low attachment 6-well plates with 4 mL of medium per well. On day 8, embryoid bodies were washed with DMEM:F12 and transferred to non-treated 6-well plates with 4 mL choroid plexus induction medium containing DMEM:F12 supplemented with 1X N2 supplement, 1X Penicillin/Streptomycin, 1X Non-essential Amino Acids, 1X GlutaMax, 3 μM CHIR-99021, and 200 ng/mL human recombinant BMP-7 per well. From this point onward, organoids were cultured on an orbital shaker rotating at 110 rpm. No more than 15 embryoid bodies were cultured in each well and half of the medium was replaced every 2 days. On day 30, medium was replaced with differentiation medium containing a 1:1 mixture of DMEM/F12 and Neurobasal supplemented with 1X N2 supplement, 1X B27 supplement w/o vitamin A, 1X Penicillin/Streptomycin, 1X Non-essential Amino Acids, 1X GlutaMax, 10 ng/mL BDNF, and 10 ng/mL GDNF. Differentiation medium was replaced every three days and organoids could be maintained for over 100 days.

### SARS-CoV-2 Infection of Cultures

SARS-CoV-2 USA-WA1/2020 (Gen Bank: MN985325.1) (Harcourt et al., 2020) viral isolate was obtained from BEI Resources. SARS-CoV-2 virus was expanded and titered using Vero E6 cells. Culture cells were infected in a 96-well plate with 100 μL of medium per well using multiplicity (MOI) of 0.2, 1, or 5, and vehicle treated cells as controls. After virus exposure for 12 hours, cells were washed 3 times with fresh medium and returned to culture. Brain organoids were infected in a 24-well ultra-low attachment plate (Corning) with 0.5 mL of medium per well containing vehicle or 10^5^ focus forming unites (FFU) of SARS-CoV-2 for 8 hours on an orbital shaker rotating at approximately 100 rpm. After virus exposure, organoids were washed 3 times with fresh medium and returned to culture.

### Tissue Processing and Immunohistology

At experimental endpoints, brain organoids were placed directly in 4% methanol-free formaldehyde (Polysciences) diluted in DPBS (Thermo Fisher Scientific) overnight at 4°C on a rocking shaker. After fixation, the brain organoids were washed in DPBS and cryoprotected by overnight incubation in 30% sucrose (Sigma-Aldrich) in DPBS at 4°C. Brain organoids were placed in a plastic cryomold (Electron Microscopy Sciences) and snap frozen in tissue freezing medium (General Data) on dry ice. Frozen tissue was stored at −80°C until processing. Monolayer cells grown on glass coverslips were fixed in 4% methanol-free formaldehyde (Polysciences), diluted in DPBS (ThermoFisher Scientific) for 5 hours at room temperature, washed with DPBS, and stored at 4°C until ready for immunohistology. Serial tissue sections (25 μm for brain organoids) were sliced using a cryostat (Leica, CM3050S), and melted onto charged slides (Thermo Fisher Scientific). Slides were dried at room temperature and stored at −20°C until ready for immunohistology. For immunofluorescence staining, the tissue sections were outlined with a hydrophobic pen (Vector Laboratories) and washed with TBS containing 0.1% Tween-20 (v/v). Tissue sections were permeabilized and non-specific binding was blocked using a solution containing 10% donkey serum (v/v), 0.5% Triton X-100 (v/v), 1% BSA (w/v), 0.1% gelatin (w/v), and 22.52 mg/ml glycine in TBST for 1 hour at room temperature. The tissue sections were incubated with primary antibodies diluted in TBST with 5% donkey serum (v/v) and 0.1% Triton X-100 (v/v) overnight at 4°C. After washing in TBST, the tissue sections were incubated with secondary antibodies diluted in TBST with 5% donkey serum (v/v) and 0.1% Triton X-100 (v/v) for 1.5 hours at room temperature. After washing with TBST, sections were washed with TBS to remove detergent and incubated with TrueBlack reagent (Biotium) diluted 1:20 in 70% ethanol for 1 minute to block autofluorescence. After washing with TBS, slides were mounted in mounting solution (Vector Laboratories), cover-slipped, and sealed with nail polish.

### Confocal Microscopy and Image Processing

Monolayer cultures were imaged with a Perkin-Elmer Opera Phenix high-content automatic imaging system with a 20x air objective in confocal mode. At least 9 fields were acquired per well. Images were analyzed using Columbus Image Data Storage and Analysis System. Brain organoid sections were imaged as z stacks using a Zeiss LSM 810 confocal microscope or a Zeiss LSM 710 confocal microscope (Zeiss) using a 10X, 20X, 40X, or 63X objective with Zen 2 software (Zeiss). Images were analyzed using either Imaris 7.6 or ImageJ software. Images were cropped and edited using Adobe Photoshop (Adobe) and Adobe Illustrator (Adobe).

### Viral Titer Assay

Tissue culture supernatants and lysates from SARS-CoV-2 treated CPOs were collected at 0, 24, and 72 hpi for 2 biological replicates per timepoint containing 4 organoids each. Tissue culture supernatants were stored at −80°C until titration. For lysates, organoids were washed with cold PBS, mechanically dispersed and disrupted by freeze-thaw cycles, and the clarified homogenate was stored at −80°C until titration. Viral titers were determined in Vero E6 cells using an immune focus forming unit assay. Briefly, Vero E6 cells (3 x 10^4^ cells/96-well) were inoculated with 100 μl of serially diluted organoid supernatant or lysate in a 96-well plate at 37°C and 5% CO_2_ for 20 hours. Cells were fixed with 4% methanol-free paraformaldehyde overnight at 4°C. Infected foci were labeled using immunofluorescence staining for SARS-CoV-2 nucleoprotein. SARS-CoV-2 nucleoprotein positive cells were counted in each well. Readouts were performed on the wells with the highest dilution displaying observable infection.

### RNA Isolation, Library Preparation, and Sequencing

To minimize variability due to sampling and processing, each biological replicate consisted of 3 organoids and the replicates for all experimental conditions were processed in parallel for RNA-extraction, library preparation and sequencing. At the desired experimental endpoints, organoids were homogenized in TRIzol (Thermo Fisher Scientific) using a disposable pestle and handheld mortar and stored at −80 °C until processing. RNA clean-up was performed using the RNA Clean & Concentrator kit (Zymo Research) after TRIzol phase separation according to the manufacturer’s protocol. RNA concentration and quality were assessed using a Nanodrop 2000 (Thermo Fisher Scientific).

Library preparation was performed as previously described with some minor modifications (Weng et al., 2017). About 300 ng of RNA in 3.2 μL was combined with 0.25 μL RNase inhibitor (NEB) and 1 μL CDS primer (5’-AAGCAGTGGTATCAACGCAGAGTACT30VN-3’) in an 8-well PCR tube strip, heated to 70 °C for 2 min, and immediately placed on ice. 5.55 μL RT mix, containing 2 μL of 5X SMARTScribe RT buffer (Takara), 0.5 μL of 100 mM DTT (Millipore Sigma), 0.3 μL of 200 mM MgCl_2_ (Thermo Fisher Scientific), 1 μL of 10 mM dNTPs (Takara), 1 μL of 10 μM TSO primer (5’-AAGCAGTGGTATCAACGCAGAGTACATrGrGrG-3’), 0.25 μL of RNase inhibitor (NEB), and 0.5 μL SMARTScribe reverse transcriptase (Takara) was added to the reaction. RT was performed under the following conditions: 42 °C for 90 minutes, 10 cycles of 50 °C for 2 minutes and 42 °C for 2 minutes, 70 °C for 15 minutes, and 4 °C indefinitely. For cDNA amplification, 2 μL of the RT reaction was combined with 2.5 μL of 10X Advantage 2 buffer (Takara), 2.5 μL of 2.5 mM dNTPs (Takara), 0.25 μL of 10 μM IS PCR primer (5’-AAGCAGTGGTATCAACGCAGAGT-3’), 17.25 μL nuclease free water (ThermoFisher), and 0.5 μL Advantage 2 DNA Polymerase (Takara). Thermocycling conditions were as follows: 94 °C for 3 minutes, 8 cycles of 94 °C for 15 s, 65 °C for 30 s, and 68 °C for 6 minutes, 72 °C for 10 minutes, and 4 °C indefinitely. Amplified cDNA was purified using 0.8X AMPure XP beads (Beckman Coulter), eluted in 15 μL nuclease-free water, and quantified using Qubit dsDNA HS assay kit (Thermo Fisher Scientific). cDNA was fragmented by combining 100 pg cDNA in 1 μL nuclease free water, 2X TD buffer (20 mM Tris, pH 8.0; Thermo Fisher Scientific), 10 mM MgCl_2_, and 16% PEG 8000 (MilliporeSigma), and 0.5 μL Tn5 (Lucigen). The mixture was heated to 55 °C for 12 minutes, and the reaction was terminated upon the addition of 1.25 μL of 0.2% SDS (Fisher) and incubated at room temperature for 10 minutes. Fragments were amplified by adding 16.75 μL nuclease free water (Thermo Fisher Scientific), 1 μL of 10 mM Nextera i7 primer, 1 μL of 10 mM Nextera i5 primer, and 25 μL KAPA HiFi hotstart readymix (EMSCO/FISHER). Thermocycling conditions were as follows: 72 °C for 5 minutes, 95 °C for 1 minute, 14 cycles of 95 °C for 30 s, 55 °C for 30 s, and 72 °C for 30 s, 72 °C for 1 minute, and 4 °C indefinitely. DNA was purified twice with 0.8X AMPure XP beads (Beckman Coulter) and eluted in 10 μL of 10 mM Tris, pH 8 (Thermo Fisher Scientific). Samples were quantified by qPCR (KAPA) and pooled at equal molar amounts. Final sequencing library fragment sizes were quantified by bioanalyzer (Agilent) with an average size of ~420 bp, and concentrations were determined by qPCR (KAPA). Samples were loaded at concentrations of 2.7 pM and sequenced on a NextSeq 550 (Illumina) using 1×72 bp reads to an average depth of 40 million reads per sample.

### Bioinformatics Analyses

Choroid plexus organoid raw sequencing data were demultiplexed with bcl2fastq2 v2.17.1.14 (Illumina) with adapter trimming using Trimmomatic v0.32 software (Bolger et al., 2014). Alignment was performed using STAR v2.5.2a (Dobin et al., 2013). A combined reference genome consisting of GENCODE human genome release 28 (GRCh38.p12) and Ensembl SARS-CoV-2 Wuhan-Hu-1 isolate genome (genome assembly: ASM985889v3, GCA_009858895.3; sequence: MN908947.3) was used for alignment. Multimapping and chimeric alignments were discarded, and uniquely mapped reads were quantified at the exon level and summarized to gene counts using STAR -- *quantMode GeneCounts*. All further analyses were performed in R v3.6.0. Transcript lengths were retrieved from GTF annotation files using GenomicFeatures v1.36.4 (Lawrence et al., 2013) and raw counts were converted to units of transcripts per million (TPM) using the *calculateTPM* function in scater v1.12.2 (McCarthy et al., 2017). A combined TPM normalization using both human and SARS-CoV-2 viral transcripts was used to obtain TPM values for SARS-CoV-2 viral transcripts. A human-only TPM normalization was used to obtain TPM values for human transcripts. TPM expression values for human choroid plexus markers, SARS-CoV-2 receptors, and SARS-CoV-2 viral transcripts were log_10_(TPM + 1) transformed for plotting.

Differential gene expression analysis for choroid plexus organoids between 72 hpi vehicle, 24 hpi SARS-CoV-2 infection, and 72 hpi SARS-CoV-2 infection was performed using DESeq2 v1.24.0 (Love et al., 2014). SARS-CoV-2 viral transcripts were removed for differential gene expression analysis. Variance stabilizing transformation (VST) of raw counts was performed prior to whole-transcriptome principal component analysis (PCA) using DESeq2 v1.24.0. Upregulated and downregulated genes for 24 hpi SARS-CoV-2 compared to 72 hpi vehicle, and 72 hpi SARS-CoV-2 compared to 72 hpi vehicle were filtered by adjusted p-value < 0.05. Volcano plots for 24 hpi SARS-CoV-2 and 72 hpi SARS-CoV-2 were plotted using EnhancedVolcano v1.7.8. Shared and unique signatures for upregulated and downregulated genes between 24 hpi SARS-CoV-2 and 72 hpi SARS-CoV-2 were visualized using VennDiagram v1.6.20 (Chen and Boutros, 2011). Upregulated and downregulated gene lists for each timepoint were used for gene ontology (GO) (Ashburner et al., 2000; The Gene Ontology, 2019) enrichment analysis using PANTHER v15.0 (Mi et al., 2019). For each gene list, a list of terms for GO biological process complete, GO molecular function complete, and GO cellular component complete were filtered by false discovery rate (FDR) < 0.05. Selected genes from notable GO terms were log_2_(TPM + 1) normalized and converted to row Z-scores per gene for visualization.

For SARS-CoV-2 cross-organoid comparison analysis, a list of upregulated and downregulated differentially expressed genes (DEGs) after SARS-CoV-2 infection were downloaded for adult hepatocyte organoids after 24 hpi SARS-CoV-2 (Yang et al., 2020a), intestinal organoids after 60 hpi SARS-CoV-2 in expansion (EXP) medium (Lamers et al., 2020), and intestinal organoids after 60 hpi SARS-CoV-2 in differentiation (DIF) medium (Lamers et al., 2020). Upregulated and downregulated genes were filtered by adjusted p-value < 0.05. Shared and unique signatures for upregulated and downregulated genes for the total gene set for choroid plexus organoids (24 hpi and 72 hpi SARS-CoV-2), adult hepatocyte organoids (24 hpi SARS-CoV-2), and intestinal organoids (60 hpi SARS-CoV-2 in EXP medium and 60 hpi SARS-CoV-2 in DIF medium) were visualized using VennDiagram v1.6.20.

### Quantification and Statistical Analysis

All statistical tests and sample sizes are included in the Figure Legends and text. All data are shown as mean ± SEM or mean + SD as stated in the Figure Legends. In all cases, the p values are represented as follows: ***p < 0.001, **p < 0.01, *p < 0.05, and not statistically significant when p > 0.05. In all cases, the stated “n” value is either separate wells of monolayer cultures or individual organoids with multiple independent images used to obtain data points for each. No statistical methods were used to pre-determine sample sizes. For all quantifications of immunohistology, the samples being compared were processed in parallel and imaged using the same settings and laser power for confocal microscopy. For monolayer cultures, cells were segmented by first identifying nuclei, and then cytoplasmic area. Mean cytoplasmic intensities for SARS-CoV-2 nucleoprotein immunostaining were then measured in the segmented cytoplasmic regions, and a mean cytoplasmic intensity threshold was determined to identify SARS-CoV-2 infected cells. Cell numbers were then automatically counted for SARS-CoV-2 infected DAPI^+^ cells, and total DAPI^+^ cells. For brain organoids, immuno-positive cells were quantified manually using the cell counter function in ImageJ (NIH) or Imaris (Bitplane).

### Data and Code Availability

The access number for the RNA-seq data reported in this study will be deposited in GEO. The data that support the findings of this study are available from the lead contact, Dr. Guo-li Ming upon reasonable request.

## REFERENCES

Ashburner, M., Ball, C.A., Blake, J.A., Botstein, D., Butler, H., Cherry, J.M., Davis, A.P., Dolinski, K., Dwight, S.S., Eppig, J.T., et al. (2000). Gene ontology: tool for the unification of biology. The Gene Ontology Consortium. Nat Genet 25, 25–29.

Bao, L., Deng, W., Huang, B., Gao, H., Liu, J., Ren, L., Wei, Q., Yu, P., Xu, Y., Qi, F., et al. (2020). The pathogenicity of SARS-CoV-2 in hACE2 transgenic mice. Nature.

Bardy, C., van den Hurk, M., Eames, T., Marchand, C., Hernandez, R.V., Kellogg, M., Gorris, M., Galet, B., Palomares, V., Brown, J., et al. (2015). Neuronal medium that supports basic synaptic functions and activity of human neurons in vitro. Proc Natl Acad Sci U S A 112, E2725–2734.

Beers, J., Linask, K.L., Chen, J.A., Siniscalchi, L.I., Lin, Y., Zheng, W., Rao, M., and Chen, G. (2015). A cost-effective and efficient reprogramming platform for large-scale production of integration-free human induced pluripotent stem cells in chemically defined culture. Scientific reports 5, 11319.

Bolger, A.M., Lohse, M., and Usadel, B. (2014). Trimmomatic: a flexible trimmer for Illumina sequence data. Bioinformatics 30, 2114–2120.

Bouhaddou, M., Memon, D., Meyer, B., White, K.M., Rezelj, V.V., Correa Marrero, M., Polacco, B.J., Melnyk, J.E., Ulferts, S., Kaake, R.M., et al. (2020). The Global Phosphorylation Landscape of SARS-CoV-2 Infection. Cell.

Brown, P.D., Davies, S.L., Speake, T., and Millar, I.D. (2004). Molecular mechanisms of cerebrospinal fluid production. Neuroscience 129, 957–970.

Cantuti-Castelvetri, L., Ojha, R., Pedro, L.D., Djannatian, M., Franz, J., Kuivanen, S., Kallio, K., Kaya, T., Anastasina, M., Smura, T., et al. (2020). Neuropilin-1 facilitates SARS-CoV-2 cell entry and provides a possible pathway into the central nervous system. BioRxiv.

Chen, H., and Boutros, P.C. (2011). VennDiagram: a package for the generation of highly-customizable Venn and Euler diagrams in R. BMC Bioinformatics 12, 35.

Chen, R., Wang, K., Yu, J., Howard, D., French, L., Chen, Z., Wen, C., and Xu, Z. (2020). The spatial and cell-type distribution of SARS-CoV-2 receptor ACE2 in human and mouse brain. Biorxiv.

Chiang, C.H., Su, Y., Wen, Z., Yoritomo, N., Ross, C.A., Margolis, R.L., Song, H., and Ming, G.L. (2011). Integration-free induced pluripotent stem cells derived from schizophrenia patients with a DISC1 mutation. Mol Psychiatry 16, 358–360.

Clevers, H. (2016). Modeling Development and Disease with Organoids. Cell 165, 1586–1597.

Desforges, M., Le Coupanec, A., Stodola, J.K., Meessen-Pinard, M., and Talbot, P.J. (2014). Human coronaviruses: viral and cellular factors involved in neuroinvasiveness and neuropathogenesis. Virus Res 194, 145–158.

Dobin, A., Davis, C.A., Schlesinger, F., Drenkow, J., Zaleski, C., Jha, S., Batut, P., Chaisson, M., and Gingeras, T.R. (2013). STAR: ultrafast universal RNA-seq aligner. Bioinformatics 29, 15–21.

Dong, L., Tian, J., He, S., Zhu, C., Wang, J., Liu, C., and Yang, J. (2020). Possible Vertical Transmission of SARS-COV-2 From an Infected Mother to Her Newborn. JAMA 323, 1846–1848.

Dube, M., Le Coupanec, A., Wong, A.H.M., Rini, J.M., Desforges, M., and Talbot, P.J. (2018). Axonal Transport Enables Neuron-to-Neuron Propagation of Human Coronavirus OC43. J Virol 92.

Ellul, M.A., Benjamin, L., Singh, B., Lant, S., Michael, B.D., Easton, A., Kneen, R., Defres, S., Sejvar, J., and Solomon, T. (2020). Neurological associations of COVID-19. Lancet Neurol.

Engelhardt, B., and Tietz, S. (2015). Brain barriers: Crosstalk between complex tight junctions and adherens junctions. Journal of Cell Biology 209, 493–506.

Farhadian, S., Glick, L.R., Vogels, C.B.F., Thomas, J., Chiarella, J., Casanovas-Massana, A., Zhou, J., Odio, C., Vijayakumar, P., Geng, B., et al. (2020). Acute encephalopathy with elevated CSF inflammatory markers as the initial presentation of COVID-19. BMC neurology 20, 248.

Ghersi-Egea, J.F., Strazielle, N., Catala, M., Silva-Vargas, V., Doetsch, F., and Engelhardt, B. (2018). Molecular anatomy and functions of the choroidal blood-cerebrospinal fluid barrier in health and disease. Acta Neuropathol 135, 337–361.

Giorgianni, A., Vinacci, G., Agosti, E., Cariddi, L.P., Mauri, M., Baruzzi, F., and Versino, M. (2020). Transient acute-onset tetraparesis in a COVID-19 patient. Spinal Cord.

Harcourt, J., Tamin, A., Lu, X., Kamili, S., Sakthivel, S.K., Murray, J., Queen, K., Tao, Y., Paden, C.R., Zhang, J., et al. (2020). Severe Acute Respiratory Syndrome Coronavirus 2 from Patient with Coronavirus Disease, United States. Emerg Infect Dis 26, 1266–1273.

Helms, J., Kremer, S., Merdji, H., Clere-Jehl, R., Schenck, M., Kummerlen, C., Collange, O., Boulay, C., Fafi-Kremer, S., Ohana, M., et al. (2020). Neurologic Features in Severe SARS-CoV-2 Infection. N Engl J Med 382, 2268–2270.

Hladky, S.B., and Barrand, M.A. (2016). Fluid and ion transfer across the blood-brain and blood-cerebrospinal fluid barriers; a comparative account of mechanisms and roles. Fluids and barriers of the CNS 13, 19.

Hoffmann, M., Kleine-Weber, H., and Pohlmann, S. (2020a). A Multibasic Cleavage Site in the Spike Protein of SARS-CoV-2 Is Essential for Infection of Human Lung Cells. Mol Cell 78, 779–784 e775.

Hoffmann, M., Kleine-Weber, H., Schroeder, S., Kruger, N., Herrler, T., Erichsen, S., Schiergens, T.S., Herrler, G., Wu, N.H., Nitsche, A., et al. (2020b). SARS-CoV-2 Cell Entry Depends on ACE2 and TMPRSS2 and Is Blocked by a Clinically Proven Protease Inhibitor. Cell 181, 271–280 e278.

Huang, J.T., Leweke, F.M., Oxley, D., Wang, L., Harris, N., Koethe, D., Gerth, C.W., Nolden, B.M., Gross, S., Schreiber, D., et al. (2006). Disease biomarkers in cerebrospinal fluid of patients with first-onset psychosis. PLoS medicine 3, e428.

Hung, E.C., Chim, S.S., Chan, P.K., Tong, Y.K., Ng, E.K., Chiu, R.W., Leung, C.B., Sung, J.J., Tam, J.S., and Lo, Y.M. (2003). Detection of SARS coronavirus RNA in the cerebrospinal fluid of a patient with severe acute respiratory syndrome. Clinical chemistry 49, 2108–2109.

Lamers, M.M., Beumer, J., van der Vaart, J., Knoops, K., Puschhof, J., Breugem, T.I., Ravelli, R.B.G., Paul van Schayck, J., Mykytyn, A.Z., Duimel, H.Q., et al. (2020). SARS-CoV-2 productively infects human gut enterocytes. Science 369, 50–54.

Lau, K.K., Yu, W.C., Chu, C.M., Lau, S.T., Sheng, B., and Yuen, K.Y. (2004). Possible central nervous system infection by SARS coronavirus. Emerg Infect Dis 10, 342–344.

Lawrence, M., Huber, W., Pages, H., Aboyoun, P., Carlson, M., Gentleman, R., Morgan, M.T., and Carey, V.J. (2013). Software for computing and annotating genomic ranges. PLoS Comput Biol 9, e1003118.

Love, M.I., Huber, W., and Anders, S. (2014). Moderated estimation of fold change and dispersion for RNA-seq data with DESeq2. Genome Biol 15, 550.

Lun, M.P., Monuki, E.S., and Lehtinen, M.K. (2015). Development and functions of the choroid plexus-cerebrospinal fluid system. Nat Rev Neurosci 16, 445–457.

Ma-Lauer, Y., Carbajo-Lozoya, J., Hein, M.Y., Muller, M.A., Deng, W., Lei, J., Meyer, B., Kusov, Y., von Brunn, B., Bairad, D.R., et al. (2016). p53 down-regulates SARS coronavirus replication and is targeted by the SARS-unique domain and PLpro via E3 ubiquitin ligase RCHY1. Proc Natl Acad Sci U S A 113, E5192–5201.

Mao, L., Jin, H., Wang, M., Hu, Y., Chen, S., He, Q., Chang, J., Hong, C., Zhou, Y., Wang, D., et al. (2020). Neurologic Manifestations of Hospitalized Patients With Coronavirus Disease 2019 in Wuhan, China. JAMA neurology.

Matsuyama, S., Nao, N., Shirato, K., Kawase, M., Saito, S., Takayama, I., Nagata, N., Sekizuka, T., Katoh, H., Kato, F., et al. (2020). Enhanced isolation of SARS-CoV-2 by TMPRSS2-expressing cells. Proc Natl Acad Sci U S A 117, 7001–7003.

McCarthy, D.J., Campbell, K.R., Lun, A.T., and Wills, Q.F. (2017). Scater: pre-processing, quality control, normalization and visualization of single-cell RNA-seq data in R. Bioinformatics 33, 1179–1186.

Mi, H., Muruganujan, A., Ebert, D., Huang, X., and Thomas, P.D. (2019). PANTHER version 14: more genomes, a new PANTHER GO-slim and improvements in enrichment analysis tools. Nucleic Acids Res 47, D419–D426.

Ming, G.L., Tang, H., and Song, H. (2016). Advances in Zika Virus Research: Stem Cell Models, Challenges, and Opportunities. Cell Stem Cell 19, 690–702.

Monteil, V., Kwon, H., Prado, P., Hagelkruys, A., Wimmer, R.A., Stahl, M., Leopoldi, A., Garreta, E., Hurtado Del Pozo, C., Prosper, F., et al. (2020). Inhibition of SARS-CoV-2 Infections in Engineered Human Tissues Using Clinical-Grade Soluble Human ACE2. Cell 181, 905–913 e907.

Moriguchi, T., Harii, N., Goto, J., Harada, D., Sugawara, H., Takamino, J., Ueno, M., Sakata, H., Kondo, K., Myose, N., et al. (2020). A first case of meningitis/encephalitis associated with SARS-Coronavirus-2. Int J Infect Dis 94, 55–58.

Murphy, O.C., Merwick, A., O’Mahony, O., Ryan, A.M., and McNamara, B. (2018). Familial Hemiplegic Migraine With Asymmetric Encephalopathy Secondary to ATP1A2 Mutation: A Case Series. Journal of clinical neurophysiology : official publication of the American Electroencephalographic Society 35, e3–e7.

Netland, J., Meyerholz, D.K., Moore, S., Cassell, M., and Perlman, S. (2008). Severe acute respiratory syndrome coronavirus infection causes neuronal death in the absence of encephalitis in mice transgenic for human ACE2. J Virol 82, 7264–7275.

Ou, X., Liu, Y., Lei, X., Li, P., Mi, D., Ren, L., Guo, L., Guo, R., Chen, T., Hu, J., et al. (2020). Characterization of spike glycoprotein of SARS-CoV-2 on virus entry and its immune cross-reactivity with SARS-CoV. Nat Commun 11, 1620.

Pellegrini, L., Bonfio, C., Chadwick, J., Begum, F., Skehel, M., and Lancaster, M.A. (2020). Human CNS barrier-forming organoids with cerebrospinal fluid production. Science.

Puelles, V.G., Lutgehetmann, M., Lindenmeyer, M.T., Sperhake, J.P., Wong, M.N., Allweiss, L., Chilla, S., Heinemann, A., Wanner, N., Liu, S., et al. (2020). Multiorgan and Renal Tropism of SARS-CoV-2. N Engl J Med.

Qian, X., Jacob, F., Song, M.M., Nguyen, H.N., Song, H., and Ming, G.L. (2018). Generation of human brain region-specific organoids using a miniaturized spinning bioreactor. Nat Protoc 13, 565–580.

Qian, X., Nguyen, H.N., Song, M.M., Hadiono, C., Ogden, S.C., Hammack, C., Yao, B., Hamersky, G.R., Jacob, F., Zhong, C., et al. (2016). Brain-Region-Specific Organoids Using Mini-bioreactors for Modeling ZIKV Exposure. Cell 165, 1238–1254.

Renu, K., Prasanna, P.L., and Valsala Gopalakrishnan, A. (2020). Coronaviruses pathogenesis, comorbidities and multi-organ damage - A review. Life sciences 255, 117839.

Rodriguez-Lorenzo, S., Ferreira Francisco, D.M., Vos, R., van Het Hof, B., Rijnsburger, M., Schroten, H., Ishikawa, H., Beaino, W., Bruggmann, R., Kooij, G., et al. (2020). Altered secretory and neuroprotective function of the choroid plexus in progressive multiple sclerosis. Acta Neuropathol Commun 8, 35.

Sachs, N.A., Sawa, A., Holmes, S.E., Ross, C.A., DeLisi, L.E., and Margolis, R.L. (2005). A frameshift mutation in Disrupted in Schizophrenia 1 in an American family with schizophrenia and schizoaffective disorder. Mol Psychiatry 10, 758–764.

Sakaguchi, H., Kadoshima, T., Soen, M., Narii, N., Ishida, Y., Ohgushi, M., Takahashi, J., Eiraku, M., and Sasai, Y. (2015). Generation of functional hippocampal neurons from self-organizing human embryonic stem cell-derived dorsomedial telencephalic tissue. Nature communications 6, 8896.

Samuels, M.H. (2014). Psychiatric and cognitive manifestations of hypothyroidism. Current opinion in endocrinology, diabetes, and obesity 21, 377–383.

Shang, J., Wan, Y., Luo, C., Ye, G., Geng, Q., Auerbach, A., and Li, F. (2020). Cell entry mechanisms of SARS-CoV-2. Proc Natl Acad Sci U S A 117, 11727–11734.

Shechter, R., London, A., and Schwartz, M. (2013). Orchestrated leukocyte recruitment to immune-privileged sites: absolute barriers versus educational gates. Nature reviews Immunology 13, 206–218.

Solomon, I.H., Normandin, E., Bhattacharyya, S., Mukerji, S.S., Keller, K., Ali, A.S., Adams, G., Hornick, J.L., Padera, R.F., Jr., and Sabeti, P. (2020). Neuropathological Features of Covid-19. N Engl J Med.

Sullivan, G.M., Hatterer, J.A., Herbert, J., Chen, X., Roose, S.P., Attia, E., Mann, J.J., Marangell, L.B., Goetz, R.R., and Gorman, J.M. (1999). Low levels of transthyretin in the CSF of depressed patients. Am J Psychiatry 156, 710–715.

Sun, S.H., Chen, Q., Gu, H.J., Yang, G., Wang, Y.X., Huang, X.Y., Liu, S.S., Zhang, N.N., Li, X.F., Xiong, R., et al. (2020). A Mouse Model of SARS-CoV-2 Infection and Pathogenesis. Cell Host Microbe 28, 124–133 e124.

Tang, H., Hammack, C., Ogden, S.C., Wen, Z., Qian, X., Li, Y., Yao, B., Shin, J., Zhang, F., Lee, E.M., et al. (2016). Zika Virus Infects Human Cortical Neural Progenitors and Attenuates Their Growth. Cell Stem Cell 18, 587–590.

Tay, M.Z., Poh, C.M., Renia, L., MacAry, P.A., and Ng, L.F.P. (2020). The trinity of COVID-19: immunity, inflammation and intervention. Nature reviews Immunology 20, 363–374.

The Gene Ontology, C. (2019). The Gene Ontology Resource: 20 years and still GOing strong. Nucleic Acids Res 47, D330–D338.

Valiuddin, H., Skwirsk, B., and Paz-Arabo, P. (2020). Acute transverse myelitis associated with SARS-CoV-2: A Case-Report. Brain, Behavior, & Immunity - Health 5, 100091.

Varatharaj, A., Thomas, N., Ellul, M.A., Davies, N.W.S., Pollak, T.A., Tenorio, E.L., Sultan, M., Easton, A., Breen, G., Zandi, M., et al. (2020). Neurological and neuropsychiatric complications of COVID-19 in 153 patients: a UK-wide surveillance study. The lancet Psychiatry.

Virani, A., Rabold, E., Hanson, T., Haag, A., Elrufay, R., Cheema, T., Balaan, M., and Bhanot, N. (2020). Guillain-Barre Syndrome associated with SARS-CoV-2 infection. IDCases, e00771.

Vivanti, A.J., Vauloup-Fellous, C., Prevot, S., Zupan, V., Suffee, C., Do Cao, J., Benachi, A., and De Luca, D. (2020). Transplacental transmission of SARS-CoV-2 infection. Nature communications 11, 3572.

Vu, M., Li, R., Baskfield, A., Lu, B., Farkhondeh, A., Gorshkov, K., Motabar, O., Beers, J., Chen, G., Zou, J., et al. (2018). Neural stem cells for disease modeling and evaluation of therapeutics for Tay-Sachs disease. Orphanet J Rare Dis 13, 152.

Wang, M., Cao, R., Zhang, L., Yang, X., Liu, J., Xu, M., Shi, Z., Hu, Z., Zhong, W., and Xiao, G. (2020). Remdesivir and chloroquine effectively inhibit the recently emerged novel coronavirus (2019-nCoV) in vitro. Cell research 30, 269–271.

Watanabe, M., Kang, Y.J., Davies, L.M., Meghpara, S., Lau, K., Chung, C.Y., Kathiriya, J., Hadjantonakis, A.K., and Monuki, E.S. (2012). BMP4 sufficiency to induce choroid plexus epithelial fate from embryonic stem cell-derived neuroepithelial progenitors. J Neurosci 32, 15934–15945.

Wen, Z., Nguyen, H.N., Guo, Z., Lalli, M.A., Wang, X., Su, Y., Kim, N.S., Yoon, K.J., Shin, J., Zhang, C., et al. (2014). Synaptic dysregulation in a human iPS cell model of mental disorders. Nature 515, 414–418.

Weng, Y.L., An, R., Cassin, J., Joseph, J., Mi, R., Wang, C., Zhong, C., Jin, S.G., Pfeifer, G.P., Bellacosa, A., et al. (2017). An Intrinsic Epigenetic Barrier for Functional Axon Regeneration. Neuron 94, 337–346 e336.

WHO (2020). Coronavirus disease (COVID-19) situation dashboard. World Health Organization website https://whosprinklrcom/ Accessed April 17.

Xia, S., Liu, M., Wang, C., Xu, W., Lan, Q., Feng, S., Qi, F., Bao, L., Du, L., Liu, S., et al. (2020). Inhibition of SARS-CoV-2 (previously 2019-nCoV) infection by a highly potent pan-coronavirus fusion inhibitor targeting its spike protein that harbors a high capacity to mediate membrane fusion. Cell research 30, 343–355.

Xu, J., Zhong, S., Liu, J., Li, L., Li, Y., Wu, X., Li, Z., Deng, P., Zhang, J., Zhong, N., et al. (2005). Detection of severe acute respiratory syndrome coronavirus in the brain: potential role of the chemokine mig in pathogenesis. Clin Infect Dis 41, 1089–1096.

Xu, M., Lee, E.M., Wen, Z., Cheng, Y., Huang, W.K., Qian, X., Tcw, J., Kouznetsova, J., Ogden, S.C., Hammack, C., et al. (2016). Identification of small-molecule inhibitors of Zika virus infection and induced neural cell death via a drug repurposing screen. Nat Med 22, 1101–1107.

Yang, L., Han, Y., Nilsson-Payant, B.E., Gupta, V., Wang, P., Duan, X., Tang, X., Zhu, J., Zhao, Z., Jaffre, F., et al. (2020a). A Human Pluripotent Stem Cell-based Platform to Study SARS-CoV-2 Tropism and Model Virus Infection in Human Cells and Organoids. Cell Stem Cell 27, 125–136 e127.

Yang, X., Yu, Y., Xu, J., Shu, H., Xia, J., Liu, H., Wu, Y., Zhang, L., Yu, Z., Fang, M., et al. (2020b). Clinical course and outcomes of critically ill patients with SARS-CoV-2 pneumonia in Wuhan, China: a single-centered, retrospective, observational study. Lancet Respir Med 8, 475–481.

Ye, M., Ren, Y., and Lv, T. (2020). Encephalitis as a clinical manifestation of COVID-19. Brain, behavior, and immunity.

Yoon, K.J., Nguyen, H.N., Ursini, G., Zhang, F., Kim, N.S., Wen, Z., Makri, G., Nauen, D., Shin, J.H., Park, Y., et al. (2014). Modeling a genetic risk for schizophrenia in iPSCs and mice reveals neural stem cell deficits associated with adherens junctions and polarity. Cell Stem Cell 15, 79–91.

Zeng, H., Xu, C., Fan, J., Tang, Y., Deng, Q., Zhang, W., and Long, X. (2020). Antibodies in Infants Born to Mothers With COVID-19 Pneumonia. JAMA 323, 1848–1849.

Zhou, J., Li, C., Liu, X., Chiu, M.C., Zhao, X., Wang, D., Wei, Y., Lee, A., Zhang, A.J., Chu, H., et al. (2020). Infection of bat and human intestinal organoids by SARS-CoV-2. Nat Med 26, 1077–1083.

